# Replication is the key barrier during the dual-host adaption of mosquito-borne flaviviruses

**DOI:** 10.1101/2021.12.21.473591

**Authors:** Yanan Zhang, Dening Liang, Fei Yuan, Yiran Yan, Zuoshu Wang, Pan Liu, Qi Yu, Xing Zhang, Xiangxi Wang, Aihua Zheng

## Abstract

Mosquito-borne flaviviruses (MBFs) adapt to a dual-host transmission circle between mosquitoes and vertebrates. Dual-host affiliated insect-specific flaviviruses (dISFs), discovered from mosquitoes, are phylogenetically similar to MBFs but do not infect vertebrates. Thus, dISF-MBF chimeras could be an ideal model to study the dual-host adaption of MBFs. Using the pseudo-infectious reporter virus particle and reverse genetics systems, we found dISFs entered vertebrate cells as efficiently as the MBFs, but failed to initiate replication. Exchange of the un-translational regions (UTRs) of Donggang virus (DONV), an dISF, with those from Zika virus (ZIKV) rescued DONV replication in vertebrate cells and critical secondary RNA structures were further mapped. Essential UTR-binding host factors were screened for ZIKV replication in vertebrate cells, displaying different binding patterns. Therefore, our data demonstrate a post-entry cross-species transmission mechanism of MBFs, while UTR-host interaction is critical for dual-host adaption.

**Significance Statement:** Most viruses have a relatively narrow host range. In contrast, vector-borne flaviviruses, such as dengue virus and Zika virus, maintain their transmission cycle between arthropods and vertebrates, belonging to different phyla. How do these viruses adapt to the distinct cellular environments of two phyla? By comparing the single-host insect specific flavivirus and dual-host Zika virus, we identified three key molecular factors that determine MBF host tropism. This study will greatly increase the understanding of entry, replication, and cross-species evolution of mosquito-borne flaviviruses.

## Introduction

The genus *Flavivirus* contains arthropod-borne flaviviruses, insect-specific flaviviruses (ISFs) and vertebrate-specific flaviviruses (also known as No Known Vector, NKVs) (1). All the known pathogenic flaviviruses are arthropod-borne flaviviruses mainly transmitted by mosquitoes and ticks (2). For example, Zika virus (ZIKV), dengue virus (DENV) and West Nile virus (WNV) are mosquito-borne flaviviruses (MBFs), while tick-borne encephalitis virus and Langat virus are tick-borne flaviviruses (TBFs). Phylogenetic analysis suggests the ISFs can be divided into classical insect-specific flaviviruses (cISFs) and dual-host affiliated insect-specific flaviviruses (dISFs) (1, 3). cISFs are phylogenetically distinct from all other known flaviviruses. Interestingly, dISFs, which is apparently not monophyletic, are phylogenetically affiliated with mosquito/vertebrate flaviviruses despite their apparent insect-restricted character (1). The evolution relationships between ISFs and arthropod-borne flaviviruses remains obscure.

Up until June of 2021, forty-one ISFs had been discovered. Sixteen of them are dISFs (3–6), such as Donggang virus (DONV), Chaoyang virus (CHAOV) (7), Nounané virus (NOUV) (8) and Kampung Karu virus (KPKV) (3). They are widely distributed in every continent with the exception of Antarctica (1). Extensive studies suggest that none of these ISFs can infect vertebrates and vertebrate cells (3, 8–10). Most of the ISFs were isolated from mosquitoes and maintained by vertical transmission in nature (1, 11, 12).

Flaviviruses are plus-stranded RNA viruses, with a genome approximately 11kb in size, including a ~100 nt 5’ untranslated region (5’ UTR), a single open reading frame (ORF), and a ~400-600 nt 3’ untranslated region (3’ UTR). Special structural elements in the UTRs were identified essential for flavivirus replication, translation, and pathogenesis in mosquitoes as well as mammalian cells. The 5’ UTR contains a big stem loop A and a small stem loop B. The 3’ UTR consists of two stem loops, two dumbbells and a big conserved 3’ stem loop. Several conserved motifs were identified to be critical for the cyclization of 5’ and 3’ UTRs (13). The 5’ UTRs of dISFs and cISFs are similar in length, while the 3’UTRs of cISFs are longer than those from dISFs. In addition, dISFs have lower CpG usage than cISFs (1).

The ORF encodes three structural proteins, capsid, precursor M (prM) and E proteins, together with seven non-structural proteins (14). The precursor M is cleaved into pr+M by furin protease in the trans-Golgi network and the resulting mature virus is covered by (M+E)2 dimer (15, 16). E protein is responsible for the receptor-binding and membrane fusion, which is also the major target of neutralizing antibodies. After docking on cell surface by interaction with cellular receptors, the virions are up-taken into endosomes by clathrin-dependent endocytosis, where the low pH triggers membrane fusion (17). Although heparan sulfate, DC-SIGN, and TIM/TAM family are involved in the entry process of flaviviruses (18–20), the receptor usage of flaviviruses is largely unknown, which limits the study of the cross-species transmission of flaviviruses.

Many viruses are host-specific and only replicate in a subset of cell types. To adapt to a new host, the virus must cross a number of barriers, such as receptor binding, membrane fusion, replication, protein expression, virus assembly, secretion, and immune defense. Among these barriers, receptor recognition and interaction is the initial step that determines the host specificity. The host jump mechanism for Avian influenza viruses, for instance, is related to key mutations in the hemagglutinin protein that change receptor-binding preference.

Here, using pseudo-infectious reporter virus particles (RVPs) as an entry studying tool, we found that there is no barrier for a panel of dISFs to enter into vertebrate cells. We further confirmed with DONV that dISFs failed to replicate in the vertebrate cells. The specific interaction between the UTRs and three host factors was critical for the replication of mosquito-borne flaviviruses. Collectively, we found a cross-species transmission mechanism of flaviviruses.

## Results

### Phylogenetic and structural analysis of dISFs and MBFs

To analyze the evolution of flaviviruses, we built a phylogenetic tree (Fig. 1A) with bootstrap scores not lower than 70 for all branches except one branch of cISF. This tree shows four major phylogenetic lineages (cISF, dISF1, dISF2, and MBF). The dISF is divided into the dISF1 and dISF2 lineages, sister to MBF. We repeated our analyses by including Hepatitis C virus (HCV) and classical swine fever virus as the outgroups and the same tree topology was generated (Fig. S1). It is known that the host range of all three groups including cISFs, dISF1s, and dISF2s is restrict to arthropods, while MBFs infect both mosquitoes and vertebrates. Given their phylogeny (Fig. 1A and S1), the most parsimonious scenario is that the common ancestor of MBF expanded its host range by invading vertebrates. However, we cannot exclude the possibility that dISFs originated from a MBF ancestor that lost the ability to infect vertebrates.

**Figure 1.**
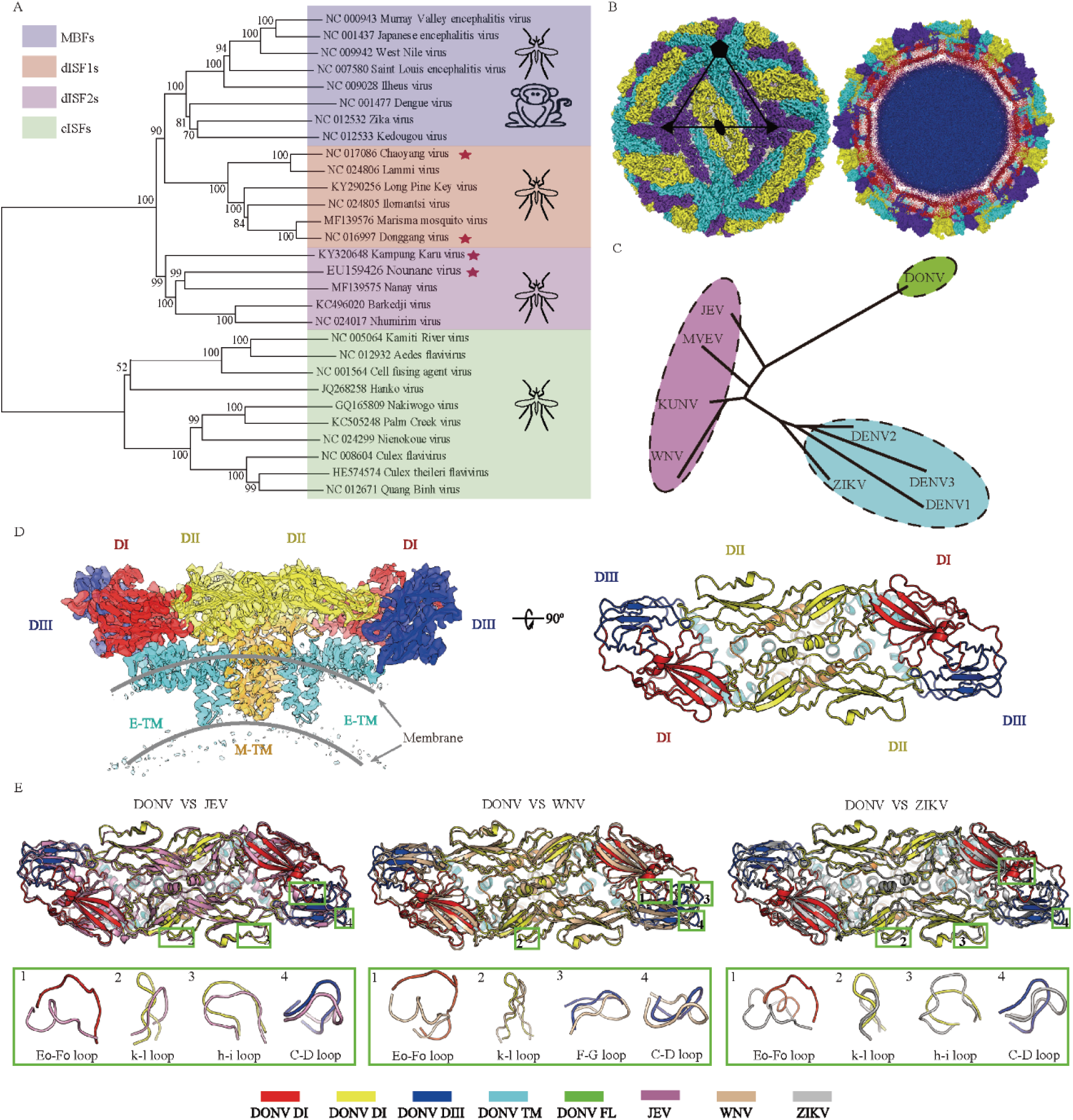
dISFs are phylogenetically and structurally close to mosquito-borne flaviviruses (MBFs). (A) Phylogenetic tree of mosquito-borne flaviviruses. Complete polyprotein amino acid sequences were aligned and a maximum likelihood phylogenetic tree was reconstructed in MEGA7. The consensus tree inferred based on 500 replicates is taken to represent the evolutionary history of the taxa. Asterisks indicate dISFs tested in Fig. 2. (B) 3D reconstruction (left) and a thin slice of the central section (right) of DONV viewed down an icosahedral twofold axis. E:M heterodimers of the same color are related by icosahedral symmetry. Heterodimers of different colors are quasi-equivalent, with cyan E:M dimers falling on the icosahedral fivefold axes, purpleon the threefold, and yellow on the twofold. (C) A structure-based phylogenetic tree constructed from E proteins of flaviviruses by Phylip-3.697: JEV, Japanese encephalitis virus (PDB code: 5WSN); KUNV, Kunjin virus (PDB code: 7KV9); MVEV, Murray Valley encephalitis virus (PDB code: 7KVB); WNV, West Nile virus (PDB code: 3J0B); ZIKV (PDB code: 6CO8); DENV 1, Dengue virus 1(PDB code: 4CCT); DENV 2, (PDB code: 7KV8); DENV 3, (PDB code: 3J6S); DONV (PDB code: 7ESD). (D) Side view of the atomic model of the E:M:M:E heterotetramer shown in ribbon. Domain I, II, III, transmembrane (TM) of E and M are colored in red, yellow, blue, cyan, and orange, respectively. (E), E and M structure superposition of DONV and JEV, WNV, ZIKV. The structures of JEV, WNV and ZIKV are shown in pink, brown and gray. The obvious differences of DONV and other flaviviruses in conformation are highlighted by the green boxes.

To further evaluate the relationship between dISFs and MBFs, the structural investigations of the Donggang virus (DONV), a dISF isolated in China (7), were performed by a cryo-electron microscope (cryo-EM). DONV was cultivated in C6/36 cells and purified using sucrose density gradient ultracentrifugation, as described previously (21, 22).Two dominant types of viral particles were observed, one exhibiting a spiky appearance with a diameter of 60 nm, characterized as immature, and the other a smooth spherical particle with a diameter of 50 nm, being a predominant form (Fig. S2). Similar to cryo-EM studies on other flaviviruses, many of either the smooth or spiky particles with local flaws (partially immature) and irregular (broken or fusogenic conformation) were observed (Fig. S2), which further confirms the theory that flaviviruses have imperfect icosahedral symmetry (23). SDS-PAGE followed by silver staining of the crude DONV particles (mixtures of spiky and smooth particles) revealed that it was mostly immature, with a prominent band for prM and only traces of mature M (Fig. S2E), akin to the experimental observation of Binjari virus (BinJV, a dISF) and some other ISFs (24), indicative of a conserved feature for ISFs. In contrast to BinJV, ~80% of the DONV particles had a smooth, rather than spiky appearance, albeit with a high level of prM at neutral environment (Figs. S2 and S3). The cryo-EM reconstruction of the spiky virion was reconstructed to a resolution of ~9 Å due to its structural flexibility (Fig. S3). 3D classification of the smooth particles showed that ~80% of the virions contained uncleaved pr, presumably due to specific subcellular location of insect furin protease (24), and less than ~20% of the smooth particles were mature virions (Fig. S3). Normal reconstruction strategies yield a reconstruction of the mature DONV particle at 4.1 Å (Fig. S4 and Table. S1). To further push the resolution, localized asymmetrical reconstruction and focused refinement by using an optimized “block-based” reconstruction strategy (25, 26) were performed. This improved the local resolution to 3.4 Å, enabling a reliable model building of the protein components of the DONV envelope, which contains 180 copies of the E and M proteins with three E and three M proteins in each icosahedral asymmetric unit (Fig. 1B and S4). The E-M-M-E heterodimers lie parallel to each other, and 90 such heterodimers cover the viral surface (Fig. 1B). A structure-based phylogenetic tree constructed from E proteins of flaviviruses indicates that DONV links the JEV group and DENV group (Fig. 1C). Like other flaviviruses, DONV E protein consists of four domains. Domains DI, DII, and DIII make up the ectodomain (Fig. 1D). The E-M-M-E heterodimers anchor to an underlying lipid bilayer envelope through their transmembrane helices E-TM and M-TM (Fig. 1D). A superposition of DONV cryo-EM structure onto the three representative (JEV, WNV, and ZIKV) cryo-EM structures reveals that these structures resemble each other (Fig. 1D). However, a number of key loops, such as E0-F0, h-i, k-l, and C-D loops that greatly differ in conformation, contribute to the serotype-specific antigenic sites (21, 27–29) and distinguish DONV from the other flaviviruses (Fig. 1E). The DONV E0-F0 loop, appearing to be the most divergent in both sequence and conformation in flaviviruses. Multiple-sequence alignment of E proteins from representative dISFs and MBFs indicates that most dISFs lack glycosylation sites at position 153/154 on E, which is distinct from the conserved glycosylated modification in MBFs (Fig. S5B) (30). In addition, the E0-F0 loops of DONV and many dISFs are enriched with over 50% charged residues (Fig. S5), including EXDDD and RKXRKEXXE motifs (Fig. S5). These distinctive structural features possibly drive viral evolution of dISF and MBF.

### dISF RVPs could enter human cells efficiently

To test whether the entry barrier restricts the infection of dISFs in vertebrate cells, we applied a reporter viral particles (RVPs) system. The RVPs were produced by trans-complementation the WNV replicon encoding a GFP reporter with the structural proteins of ISFs (31). The RVPs only support single-round infection, and the WNV replicon can replicate in both mosquito and vertebrate cells. Four out of nine dISF RVPs were successfully secreted as determined by real-time PCR to detect the WNV replicon RNAs. They were the RVPs of DONV and Chaoyang virus (CHAOV) belonging to dISF1, Nounané virus (NOUV) and Kampung Karu virus (KPKV) belonging to dISF2. Notably, the secretion efficiency of dISF RVPs was lower than the positive control ZIKV RVPs (Fig. 2A). The dISF RVPs were able to infect both mosquito cell line C6/36 and the human cell line Huh7.5, exhibiting lower titers than ZIKV as measured by infectious unit (IU) per milliliter of supernatant (Fig. 2B, 2D). The infectivity of RVPs was expressed as GE(genome)/IU. The infectivity of DONV and KPKV in Huh7.5 cells was similar to that of ZIKV, while infectivity of CHAOV and NOUV was higher than ZIKV (Fig. 2C, 2E). These results suggest that dISFs can enter human cells efficiently.

**Figure 2.**
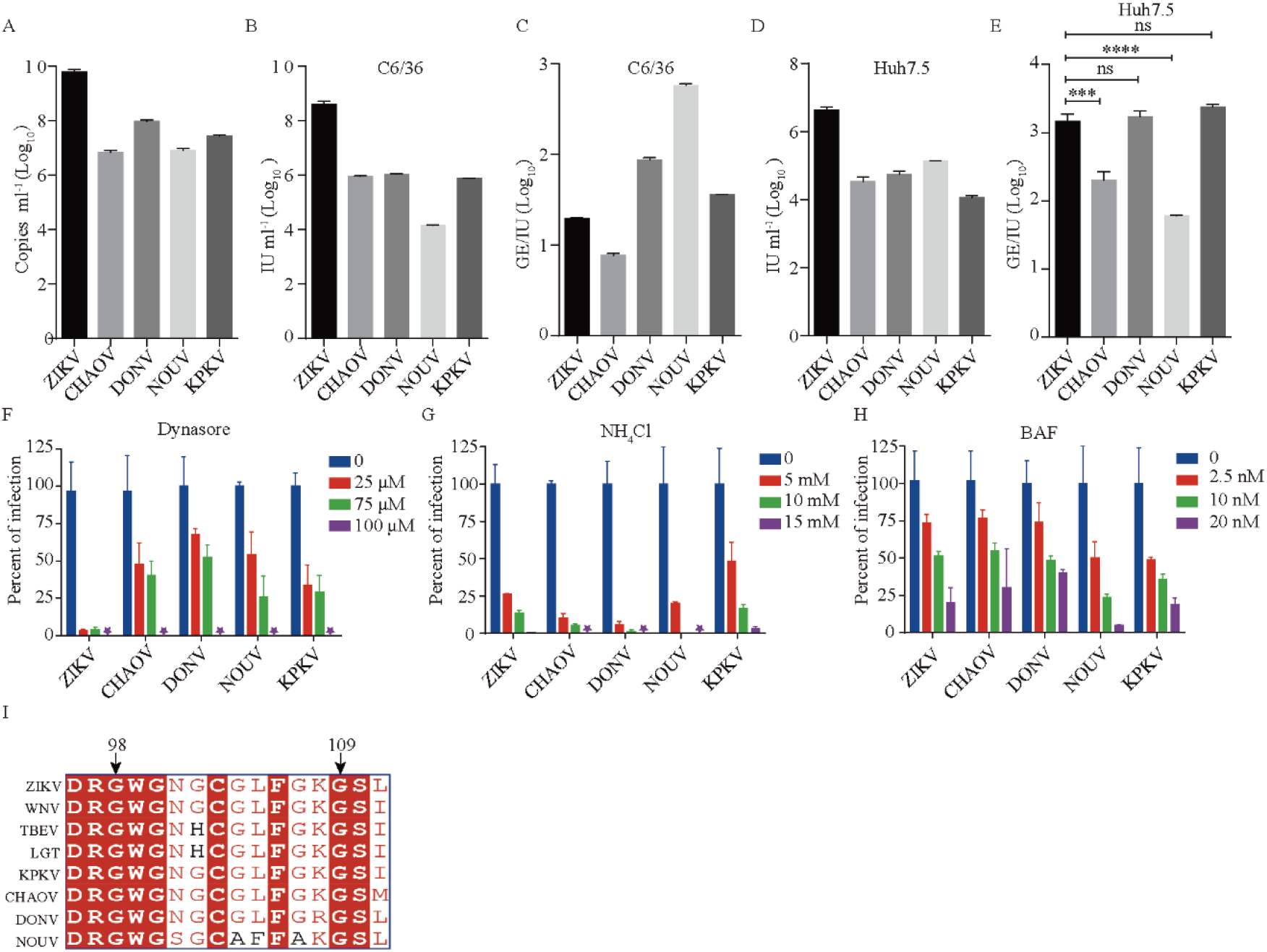
RVPs of dISFs can infect human cell. (A) RVPs were produced by co-transfection of CprME-expression plasmid with WNV replicon and RVP secretion was measured by real-time PCR using primers targeting the WNV replicon. RVP titers in the supernatant were gauged on C6/36 (B) or Huh7.5 (D) at 72 hours post-transfection by a focus-forming assay. (C and E) The infectivity of RVPs was further calculated as GE (genome)/IU. The data were analyzed by unpaired *t* test (E). (n=3). dISFs and MBFs are similar in endocytosis and membrane fusion. Huh7.5 cells were pretreated with the indicated concentrations of Dynasore (F), NH4Cl (G), or bafilomycin A (BAF) (H). One h later, RVPs of dISFs and ZIKV were added to Huh7.5 cells. The titers of RVPs were measured by a focus-forming assay 48 h post infection and the samples without compounds were set as 100%. (I) Alignment of the fusion loop of dISFs and MBFs. The titers of RVPs were measured 48 h post infection as above. Error bars indicate SD. The results are representative of three independent experiments.

Flaviviruses enter cells via clathrin-mediated endocytosis. Dynasore is an inhibitor of dynamin involved in clathrin-mediated endocytosis. Infection of both dISFs and ZIKV was reduced in the presence of dynasore in a dose-dependent manner (Fig. 2F). Membrane fusion of arthropod-borne flaviviruses is driven by low-pH and blocked by mild alkaline amino chloride (NH4Cl) and the vacuolar-type H^+^-ATPase inhibitor Bafilomycin A (BAF). The infection of dISFs was inhibited by BAF and NH4Cl in a concentration-dependent manner, similar to ZIKV (Fig. 2G, 2H). The alignment analysis of E protein fusion loop regions (residues 98–109 in ZIKV E) demonstrated a high sequence identity among dISFs and MBFs, implying a conserved membrane insertion mechanism (Fig. 2I). These results indicate that dISFs display entry characteristics similar to the MBFs.

### Authentic DONV enters endosome but cannot replicate in vertebrate cells

To further characterize the infection barrier of dISFs in vertebrate cells, a DONV infectious clone was constructed using the same strategy previously reported for ZIKV (30). The cDNA of the DONV genome was cloned into a low copy plasmid under a CMV promotor, which works in both mammalian and insect cells (32) (Fig. 3A). ZIKV could be rescued in both 293T and C6/36 cells, yet DONV only in C6/36 cells (Fig. 3B). The infectious DONV propagated efficiently in C6/36 cells, with a titer of 10^7^ FFU/ml at 48 h after infection, while no propagation was detected in Huh7.5 cells (Fig. 3C). The growth curve of DONV was also determined in C6/36 and three vertebrate cell lines including BHK-21, Vero, and Huh7.5 cells by real-time PCR. The viral RNAs were only detected in the supernatant of C6/36 cells (Fig. 3D).

**Figure 3.**
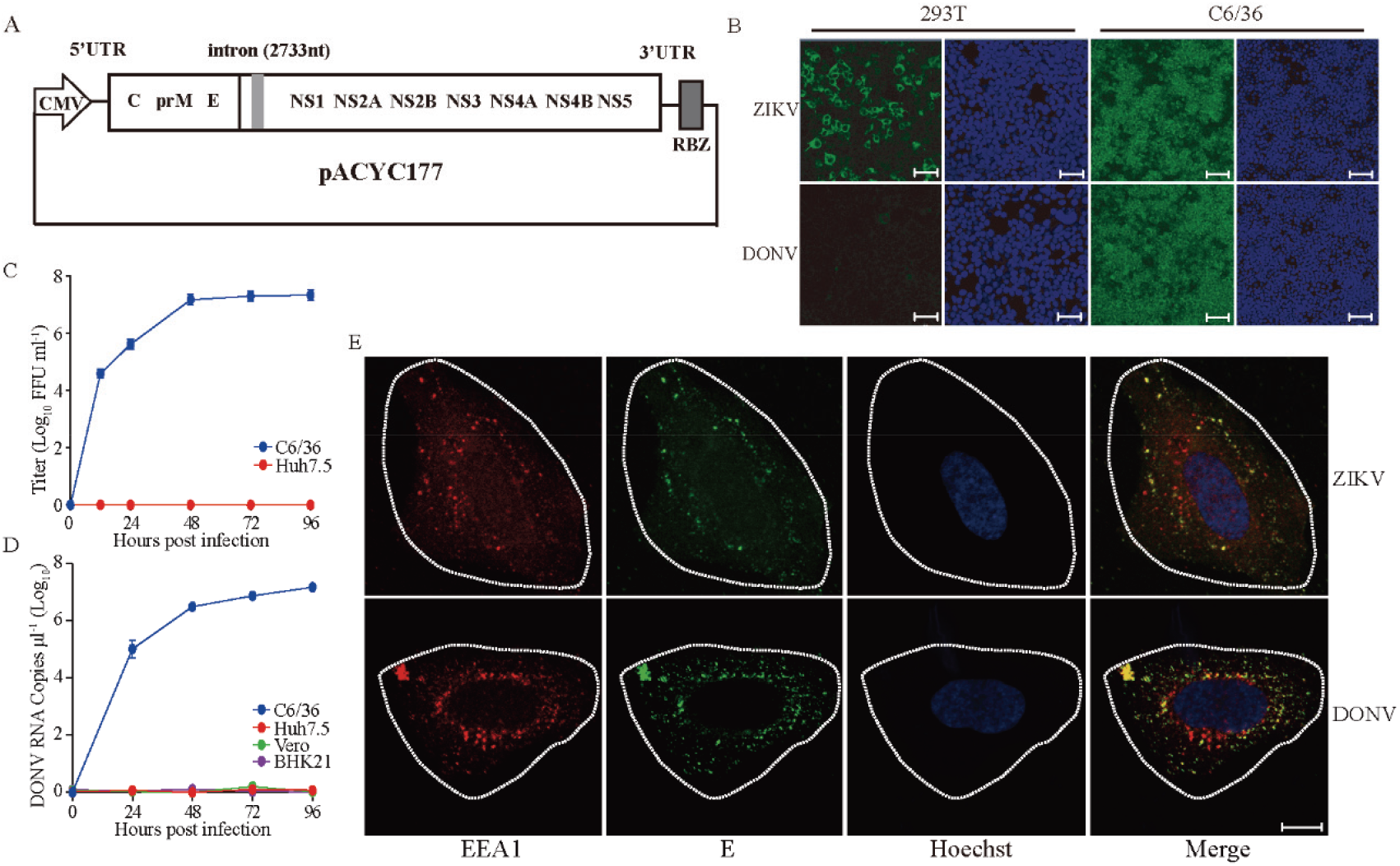
DONV can enter Vero cells but cannot replicate. (A) Schematic diagram of the DONV infectious clone. The cDNA of DONV was cloned into pACYC177 vector with a CMV promoter and an HDV RBZ termination sequence. (B) ZIKV and DONV infectious clones were transfected into 293T and C6/36 cells. Virus replication was detected by immunofluorescence with 4G2 antibody. Nuclei were stained by Hoechst 33342. Scale bar represents 100 μm. (C) Growth curve of virus on Huh7.5 and C6/36. Viral titers in the supernatant were determined at indicated time points on C6/36 cells by immunofluorescence assay. (D) C6/36, BHK21, Vero, and Huh7.5 cell lines were infected with DONV at an MOI of 0.1. Vrial RNAs in the supernatant were detected by real-time PCR at indicated time points. n=3. Error bars indicate SD. (E) Virus endocytosis was detected by colocalization of E protein and the early endosome marker EEA1 in Huh7.5 cells. E protein, EEA1, and nucleus were stained by 4G2 antibody, anti-EEA1 antibody and Hoechst 33342 respectively. Scale bar represents 10μm. These results are representative of three independent experiments.

To study the internalization of dISFs, authentic DONV and ZIKV were incubated with Huh7.5 cells. Then the cells were heated to 37°C to allow endocytosis. As shown in Fig. 3E, both DONV and ZIKV appeared in the cytoplasm of Huh7.5 cells and showed co-localization with EEA1, an early endosome marker. Combined with the data from RVPs, we demonstrated that dISFs enter vertebrate cells in a manner similar to the arthropod-borne flaviviruses, but fail to produce infectious viral particles.

### 5’ and 3’-UTRs are critical for DONV replication in vertebrate cells

The 5’ and 3’ UTRs are critical for flavivirus replication. Sequence alignment and secondary structure prediction reveals that the UTRs of DONV are different from those of ZIKV. DONV lacks a stem loop SLB in the 5’ and SL1 in the 3’ (Fig. 4A and S10). To study DONV and ZIKV replication, the viral genome copies in the cell lysates were measured after transfection of infectious clone plasmids into 293T at 37°C or C6/36 cells at 28°C. Fig. 4C shows that ZIKV RNA was detected in both 293T cells and C6/36 cells, indicating efficient replication in both vertebrate and mosquito cells. In contrast, DONV RNA was only detected in C6/36 cells, in keeping with its restricted host range. These results were consistent with double-stranded RNA (dsRNA) staining and E protein expression (Fig. 4D). As expected, the GAA replication-deficient mutants in the NS5 polymerase showed no replication (Fig. 4C and 4D). Replacement the 5’ and 3’ UTRs of DONV with those of ZIKV significantly promoted DONV RNA replication in 293T cells by 3 logs, with robust E protein expression (Fig. 4B–D). Replacement of 5’ or 3’ UTR separately (named as DONV 5’ and DONV 3’) showed the similar effects in 293T cells, irrespective of the introduction of two non-complimentary nts between the 5’ CS and 3’ CS (Fig. 4A and S7A) (33). The 5’-UTR of ZIKV consists of SLA and SLB, which are 70 nt and 30 nt in length separated by a poly (U) sequence (34, 35) (Fig. 4A and S10). However, SLA and SLB of DONV were predicted to 90 nt and 20 nt in length with no intervening poly(U) sequence, which is believed to be important for the virus replication (34). Replacing the SLA or SLB of DONV with that of ZIKV (designated as DONV 5’ 1– 81 or DONV 5’ 82–107) also enhanced RNA replication in 293T cells by real-time PCR and dsRNA staining, yet showed no expression of E protein, implying that the complete 5’ UTR was important for the viral protein translation (Fig. 4B–D).

**Figure 4.**
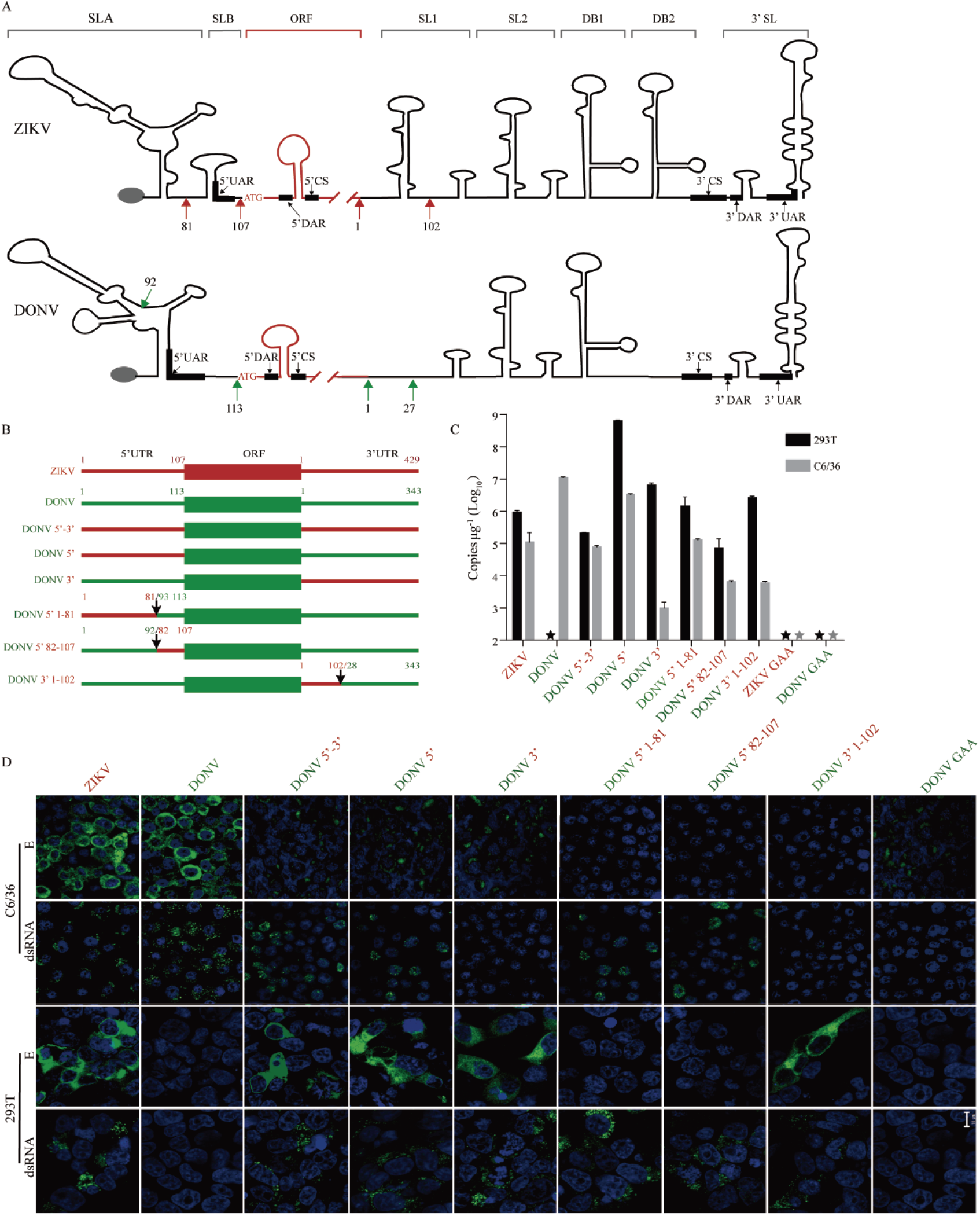
UTRs of flavivirus are determinant of dISFs replication in vertebrate cell. (A) Schematic view of secondary structure of ZIKV and DONV UTRs. (SLA, stem loop A; SLB, stem loop B; ORF, open reading frame; SL1, stem loop 1; SL2, stem loop 2; DB1, dumbbell 1; DB2, dumbbell 2; 3’SL, 3’ stem loop). Secondary structure of ZIKV and DONV UTRs were predicted by RNAfold. 5’ UAR, 5’ DAR, 5’ CS, 3’ UAR, 3’ DAR, and 3’ CS are the cyclization motifs (33). (B) Sketch map of chimeric infectious clones. Arrows represent nucleotide positions. DONV 5’-3’, the UTRs of DONV replaced by those of ZIKV, GenBank accession number, MZ357896; DONV 5’, the 5’ UTR of DONV replaced by ZIKV 5’ UTR, GenBank accession number, MZ357894; DONV 3’, the 3’ UTR of DONV replaced by ZIKV 3’ UTR, GenBank accession number, MZ357895; DONV 5’ 1-81, the 1-92nt of DONV 5’ UTR replaced by 1-81nt of ZIKV 5’ UTR, GenBank accession number, MZ357897; DONV 5’ 82-107, the 93-113 of DONV 5’UTR replaced by that of 82-107nt of ZIKV 5’UTR, GenBank accession number, MZ357898; DONV 3’ 1-102, the 1-27nt of DONV 3’UTR replaced by 1-102nt of ZIKV 3’ UTR, GenBank accession number, MZ357899. (C) Viral RNA replication in human cell 293T and insect cell C6/36 was measured by real-time PCR. n=3. Error bars indicate SD. (D) Immunofluorescence of viral double-strand RNA by anti-dsRNA mAb and envelop protein E by 4G2 mAb in transfected human cell 293T and insect cell C6/36. Nucleus was stained by Hoechst 33342. Scale bar represents 10 μm. These results are representative of three independent experiments.

The 3’-UTR of ZIKV contains duplicated SL1/SL2, DB1/DB2, and a conserved 3’SL, whereas the DONV was predicted to lack one SL (Fig. 4A). Introduction of the SL1 from ZIKV into DONV by replacing the DONV 3’ 1–27 with ZIKV 3’ 1–102 (DONV 3’ 1–102) promoted DONV RNA replication in 293T cells to a similar level as the 3’ full length replacement (Fig. 4B–C). The entire ZIKV 5’ UTR (both SLA and SLB) and/or the SL1 of ZIKV 3’ UTR appear to be important for facilitating replication of the DONV genome in vertebrate cells. However, neither replacement of the entire 3’-UTR of DONV with that of ZIKV nor the inclusion the ZIKV 3’ SL1 could maintain DONV RNA replication in C6/36 cells, implying that the 3’ UTR of DONV is critical for viral replication in insect cells. In addition, the E protein expression of DONV in C6/36 cells was obviously reduced once the UTR was altered. These data indicate that 5’ UTR and 3’ UTR, especially SL1, were critical for DONV replication in vertebrate cells.

To further confirm the role of UTRs in flavivirus host adaption, ZIKV UTRs were replaced with those of DONV. As expected, ZIKV replication in vertebrate cells was completely abolished, while only mild decrease was observed in the mosquito cells (Fig. S8). To rule out the possibility that the replacement of the UTRs could affect viral protein translation, GAA mutation was introduced into the NS5 of DONV 5’, DONV 3’, and DONV 5’-3’. As shown in Fig. S9, viral RNA replication in 293T cells was abolished, whereas E protein translation was not obviously affected. The virus release of DONV 5’, DONV 3’, and DONV 5’-3’ in 293T cells was partially rescued at 28 °C (Fig. S7), suggesting more barriers might exist in the virus assembly or release step.

### UTR-binding proteins are critical for vertebrate adaption

We speculate that the inability of DONV to replicate in vertebrate cells may be due to the failure of replication machinery formation. Using the whole 5’ and 3’ UTRs fused with an aptamer as bait, we performed a pull-down assay to identify whether vertebrate factors prefer to bind ZIKV UTRs instead of DONV UTRs (Fig. 5A). In three independent experiments, 49, 82, and 121 cytoplasmic proteins were identified with ZIKV UTR preference (Fig. 5C–E). Of the fourteen cytoplasmic proteins that appeared at least twice, three were selected for further evaluation (Fig. 5F and G). Knockout of WTAP, SYNCRIP, and G3BP1 in 293T cells significantly reduced ZIKV and DENV replication (Fig. 6A and S11A). The viral RNA in the supernatant was reduced correspondingly (Fig. 6B and S11B) and fully rescued after transduction of the corresponding genes (Fig. S12D-F). We also analyzed the binding abilities of these proteins with in vitro transcribed UTRs using an electrophoretic mobility shift assay (EMSA). Recombinant WTAP, G3BP1, and SYNCRIP proteins (Fig. S13) bound with ZIKV 5’-3’ UTRs efficiently, while no binding was detected with the DONV 5’3’ UTRs (Fig. 6C). SYNCRIP showed interaction with 5’ UTR or 3’ UTR of ZIKV, while G3BP1 only bound to the ZIKV 3’ UTR. In contrast, WTAP failed to bind 5’ or 3’ UTR of ZIKV separately (Fig. 6D and E). None of the three proteins bound the DONV UTRs. As mentioned above, the major difference in the 3’ UTR between ZIKV and DONV is lack of SL1 in DONV (Fig. 6F). We found that deletion of SL1 (1-102nt) in ZIKV 3’ UTR disrupted the SYNCRIP and G3BP1 interaction, whereas replacement of ZIKV 3’ SL1 (1-102nt) with 1-27nt of DONV 3’ UTR rescued the SYNCRIP and G3BP1 binding DONV 3’ UTR (Fig. 6G). These results suggest that WTAP, G3BP1 and SYNCRIP exhibited various binding characteristics with ZIKV UTRs and might play critical roles during MBFs replication in vertebrate cells.

**Figure 5.**
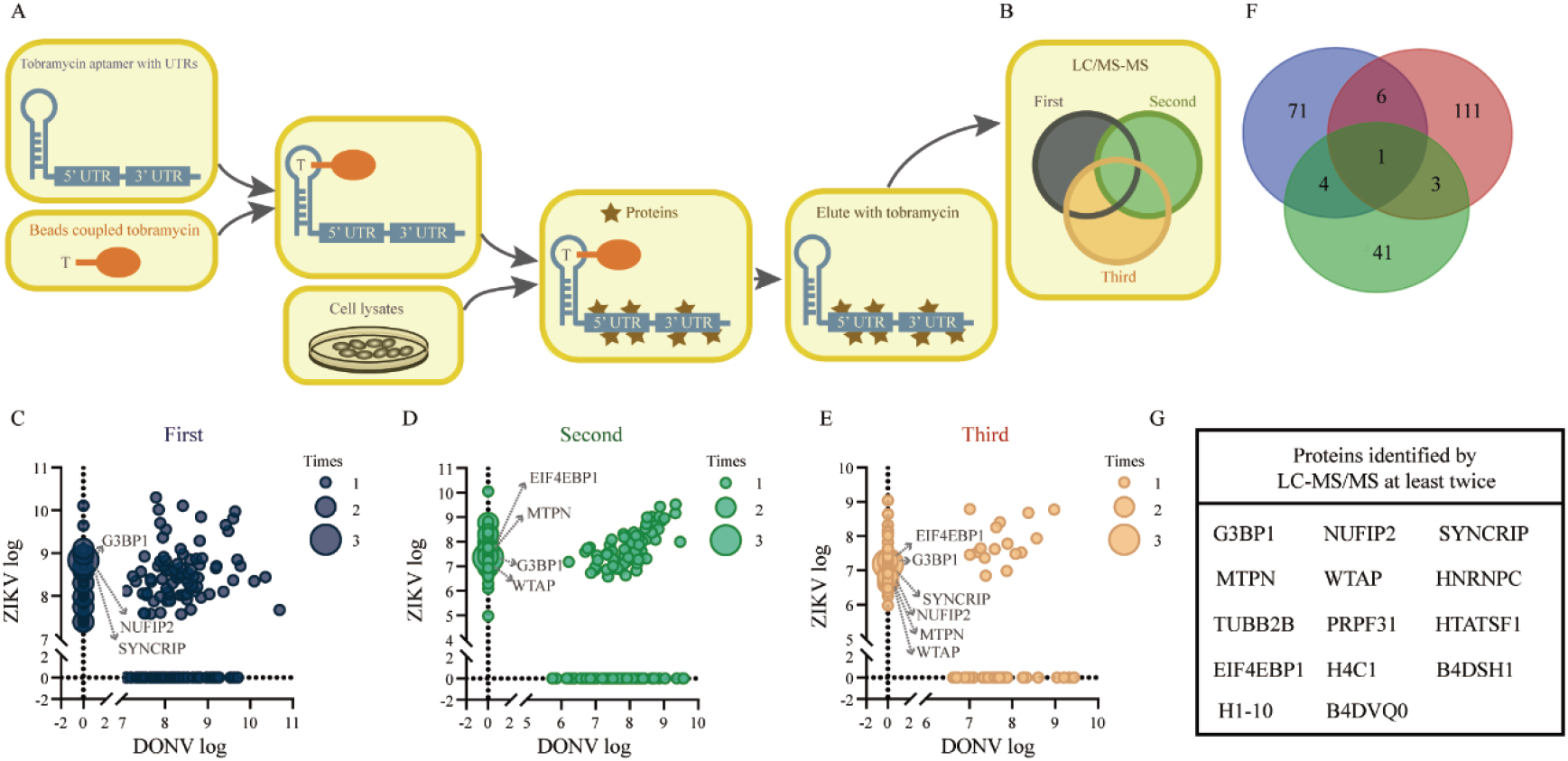
RNA affinity chromatography identified host proteins associated with ZIKV. (A and B) Outline of identification UTR-binding proteins by LC-MS/MS. (C-E) Scatter plots depicting the enrichment of the host proteins identified by LC-MS/MS with DONV and ZIKV UTRs. A total number of 238 enriched specific proteins were identified for ZIKV and 147 enriched specific proteins for DONV. (F) Venn diagram of ZIKV specific host proteins identified by LC-MS/MS. (G) List of proteins identified by LC-MS/MS at least twice.

**Figure 6.**
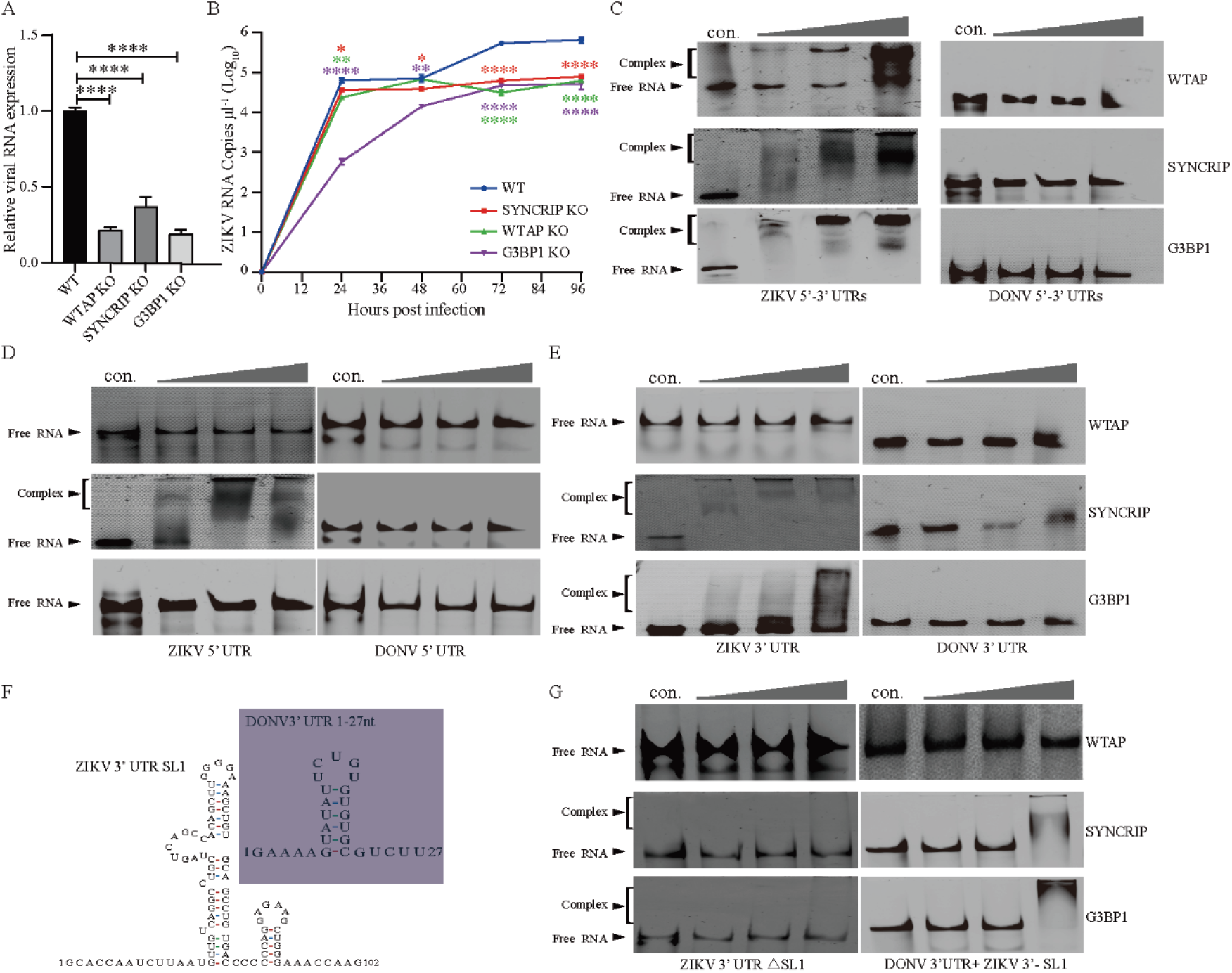
SYNCRIP, WTAP and G3BP1 have impact for ZIKV replication and bind with UTR at different positions. (A) ZIKV replication in WT, WTAP-KO (WTAP-knockout), G3BP1-KO, and SYNCRIP-KO 293T cells. The cells were infected with ZIKV at an MOI of 0.1 and the RNAs were isolated from cells 72h post infection. RNA levels were measured by real-time PCR and normalized to relative to β-actin (WT was set as 1). (B) ZIKV virus in the supernatants in above cells at indicated time were tested by real-time PCR. A and B, the data was analyzed by unpaired *t* test (A) and multiple *t* test (B). n=3. Error bars indicate SD. (C-G), The binding of WTAP, SYNCRIP, and G3BP1 with UTRs were tested by EMSA. Synthesized UTRs were incubated with indicated recombinant proteins at concentrations of 0, 0.125, 0.25, and 0.5 μg/μl (from left to right) for 30 min, then separated on native PAGE gels. (F) Predicted secondary structure of ZIKV 3’-SL1 and DONV 3’ 1-27nt. (G) ZIKV 3’ UTR with SL1 deletion (ZIKV 3’ UTR-ΔSL1) and DONV 3’UTR with 1-27nt replaced with SL1 (1-102nt) of ZIKV 3’ UTR (DONV 3’UTR + ZIKV SL1) interaction with indicated proteins. Above results are representative of three independent experiments.

## Discussion

Mosquito-borne flaviviruses have adapted to the distinct environments of mosquitoes and vertebrates. We expect that receptor adaption would be a critical barrier for dISFs in vertebrates because the receptor-virus interaction involves precise and specific structural complementation as revealed for other virus (36). However, we found a panel of dISFs could enter vertebrate cells efficiently and replication was the key barrier for dual-host adaption. Our results indicate that a specific secondary structure of the UTR is required to interact with specific host factors to initiate replication during vertebrate host invasion.

Metagenomics studies on invertebrates have discovered many insect-specific, arbovirus-like viruses (37). The high genomic complexity and diversity suggests that invertebrates are the center for the evolution of flavivirus and ISFs might be the ancestors of arthropod-borne flaviviruses (38). A number of studies have been performed to investigate the cross-species barriers of ISFs in vertebrate cells. The cISFs such as Niénokoué virus (NIEV) and Parramatta River virus are shown to have entry, replication, assembly, and release barriers in vertebrates (39, 40). Insertion of the yellow fever virus 3’UTR downstream of the NIEV ORF stop codon can enhance the viral protein translation in BHK cells but the chimera is still unable to replicate (40). Infection of two dISFs, BinJV and Aripo virus (ARPV), in vertebrates is mainly restricted in the post-entry step, which is likely mediated by the innate immune response or temperature (5, 41). The failure of Long Pine Key virus, a newly identified dISF, to infect vertebrate cells is probably due to entry and post-translational restrictions (6). The genomes of ISFs contain more CpG dinucleotides compared to vector-borne flaviviruses and the zinc-finger antiviral protein in vertebrates can bind CpG dinucleotides in the viral RNA to limit virus replication(42, 43). Here, we found dISFs are genetically and structurally close to the MBFs as revealed by the cryo-EM structure of DONV. Thus, dISFs are useful for studying the dual-host adaption of MBF. Sequence analysis suggests that the genome lengths of 5’ UTR and 3’ UTR of dISFs and MBFs are similar (1). However, dISFs could not replicate in vertebrates, although entry is still efficient. The efficient entry of dISFs gives them the chance to adapt to the environment of vertebrate cells by obtaining continuous mutations.

UTRs are important for flavivirus replication, assembly, and immune modulation. The subgenomic flaviviral RNAs (sfRNAs) are derived from the 3’UTR by incomplete degradation of the genomic RNA by XRN-1 (44). The sfRNAs are important for flavivirus replication by antagonizing the interferon pathway. The duplication of SL1 and SL2 in the DENV 3’UTR also plays critical roles in the dual-host tropism of DENV (45). The 3’ SL in the DENV 3’UTR is important for virus replication and host switching between mosquitos and vertebrates (46). Here, we found both the 5’ and 3’ UTRs are involved in the dual-host adaption of MBFs. Replacement of the whole 5’UTR with that of ZIKV greatly increased DONV replication in vertebrate cells, while changes in the SLA or SLB only resulted in a mild replication enhancement and failure of E protein expression. These results suggest that SLA and SLB might function cooperatively during both replication and translation. Replacement of the whole 3’UTR from ZIKV in DONV showed a similar phenotype as the SL1 replacement, suggesting that SL1 is the critical element involved in host adaption. However, the E protein expression of DONV in the C6/36 cells was significantly decreased by the replacement of the 5’ or/and 3’ UTRs of DONV with those of ZIKV demonstrating a critical role of UTRs in viral translation. Temperature adaption is also important for flaviviruses due to the alternative replication between vertebrates and mosquitos. Low temperature culture of 28°C is preferred by several ISFs (5, 47, 48). Similarly, the virus release of DONV 5’, DONV 3’ and DONV 5’-3’ in vertebrate cells was partially rescued only at 28°C

By pull-down with UTRs and functional analysis, we identified three host factors involved in ZIKV replication. G3BP1 is localized in the stress granules and binds the variable region stem loops in the 3’ UTR of DENV (49). These results indicate that G3BP1 preferentially bound the SL1 in the 3’ UTR of ZIKV. WTAP is a nuclear protein involved in RNA splicing with a small portion localized in the plasma (50). No reports suggest that WTAP is involved in the life cycle of any virus. WTAP only bound the linked 5’-3’ UTRs, with no interaction detected with either 5’ or 3’ UTR, suggesting it might be involved in the cyclization of ZIKV viral RNA. SYNCRIP is reported to bind HCV RNA during the replication step (51). During ZIKV replication, it interacted with both 5’ and 3’ UTRs, and shared the same binding site with G3BP1 on the SL1. All three host factors failed to bind with DONV UTRs. RNA structure prediction revealed that DONV lacks the SL1 in the 3’ UTR as compared with MBFs, such as DENV and ZIKV. Introduction of ZIKV SL1 into DONV rescued its interaction with G3BP1 and SYNCRIP, and thereby the replication in vertebrate cells. Thus, it appears that MBFs need to acquire specific RNA structures during the dual-host adaption to successfully interact with the host factors. However, DONV viral particle release in vertebrate cells was partially rescued to levels obviously lower than ZIKV after the replacement of UTRs at 28°C. It’s plausible that elements involved in virus assembly or release might be destroyed due to the replacement of UTRs. Thus, more cross-species barriers during viral assembly and secretion need to be further investigated.

## Materials and Methods

### Cell lines and reagents

C6/36, Aag2, Huh7.5, Hela, 293T, and Vero cells were originally obtained from the ATCC and were maintained as previously described (30). Huh7.5, Hela, and 293T and their KO derivatives were cultured in DMEM media supplemented with 10% FBS, 1% penicillin-streptomycin, and 1% l-glutamine at 37°C with 5% CO_2_. C6/36 cells were cultured in RPMI media supplemented with 10% FBS, 1% penicillin-streptomycin, and 1% L-glutamine at 28°C with 5% CO_2_.

A CRISPR–Cas9 strategy was employed to generate WTAP, SYNCRIP, and G3BP1-KO 293T cell lines. CRISPR guide RNA sequences were designed using a CRISPR design tool (https://zlab.bio/guide-design-resources), and the corresponding oligos were synthesized from TSINGKE Biological Technology. The oligos were cloned into the Cas9-expressing pX330 guide RNA vector. The cloning products were transfected into 293T cells using Lipofectamine 2000 (Thermo Fisher Scientific, USA) and subsequently single-cell sorted based on GFP expression into 96-well plates using a BD Influx cell sorter. Clonal cell lines were cultured to expand from a single cell, and genomic DNA was isolated for PCR-based genotyping and determined by Sanger sequencing of targeted genes.

To transduce WTAP, SYNCRIP or G3BP1 into KO cells, lentiviruses were packaged by transfecting 293T cells with 12.5 μg lentivirus plasmids carrying indicated cDNAs, 7.5 μg psPAX2 and 5 μg pMD2.G using calcium phosphate method. The supernatant was collected 48 h post-transfection and added to the target cells. After infection for 48 h, the cells were selected with 3 μg/ml puromycin for 7 d.

### Viruses, antibodies, and infectious clone

Donggang virus (DONV) was isolated from wild mosquitoes in Donggang, Liaoning Province and kept in our lab (7). 4G2 is a mouse monoclonal antibody recognizing the fusion loop of flaviviruses (52). Rabbit anti-human EEA1 monoclonal antibody conjugated with Alexa Fluor 594 label (ab206913) was purchased from Abcam (UK). DONV infectious clone was synthesized by SYKMGENE Beijing using NC_016997.1 (GenBank accession number) as the template and divided into three pieces. All three pieces were assembled into a pACYC177 vector with a CMV (Human cytomegalovirus) promotor at the 5’ end and a hepatitis delta virus (HDV) ribozyme (RBZ) terminal site at the 3’ terminus as previously (30).

DONV was rescued by transfection into C6/36 cells by FuGENE 6 transfection reagent. The virus was collected 3 d post-transfection and stored at −80°C. The titer was measured in C6/36 cells by immunofluorescence assay using 4G2 antibody.

### Particle production and purification

After removing the cell debris, the virus supernatant was ultra-filtered using a 0.22 μm filter, concentrated with a 300 kD cutoff concentrator, and then subjected to a 20–45% (w/v) discontinuous sucrose gradient ultracentrifugation at 28,000 rpm for 4 h in a P28S rotor (HIMAC, Japan) at 4°C. DONV concentrate (~ 0.3 mg in phosphate-buffered saline (PBS) buffer pH7.4) was loaded onto a 15–45% (w/v) sucrose density gradient and centrifuged at 30,000 rpm for 3.5 h in an SW40 rotor (HIMAC, Japan) at 4°C. The purified virions were determined by SDS-PAGE followed by silver staining. Fractions containing DONV were collected and dialyzed against PBS buffer. The DONV sample was concentrated to 4 mg/ml for later use.

### Cryo-EM data collection

A 3.5 μl aliquot of purified DONV virions (4 mg/ml) was applied to a freshly glow-discharged 200-mesh holey carbon-coated copper grid (C-flat, CF-2/1-2C; Protochips). Grids were blotted for 3 s in 100% relative humidity for plunge freezing (Vitrobot; FEI) in liquid ethane. Cryo-EM data sets were collected at 300 kV with an FEI Tecnai G2 Polara microscope (FEI, Hillsboro, OR, USA), equipped with a direct electron detector (K2 Summit; Gatan, Pleasanton, CA, USA). Movies (30 frames, each 0.2 s, total dose 35 e^-^ Å^-2^) were recorded with a defocus between 1.5 and 2.5 μm in single electron counting mode using SerialEM (53) at a calibrated magnification of 59,000 ×. This resulted in a pixel size of 1.35 Å.

### Image analysis, model building, and refinement

Micrographs were corrected for beam-induced drift using MOTIONCORR (54). A total of 70,022 good particles were selected by visual inspection from 2099 cryo-EM micrographs. Particles were picked automatically using the Laplacian in RELION3 (55). Contrast transfer function (CTF) parameters for each particle were estimated using Gctf47. Micrographs with signs of astigmatism or significant drift were discarded. The structure was determined using RELION3 (55) with icosahedral symmetry applied. Two-dimensional (2D) alignment was performed in RELION3 (55).

The initial model for 3D classification and refinement was generated by 3D-initial-model in RELION3 (55). A total of 3,864 particles were used to obtain the final density map at 4.1 Å, as evaluated by Fourier shell correction (threshold = 0.143 criterion), using gold-standard refinement. To improve the overall resolution, we used a block-based reconstruction strategy (56) for focusing classification and refinement. The orientation parameters of each particle determined in Relion3 were used to guide extraction of the block region, and these blocks were further 3D classified, refined, and post-processed, yielding a resolution of 3.4 Å. The atomic model of ZIKV (PDB code: 6CO8) was initially fitted into our map with CHIMERA (57) and future corrected manually by real-space refinement in COOT. This model was further refined by positional and B-factor refinement in real space with Phenix (58).

### Reporter viral particles

Human codon-optimized sequences encoding the CprME proteins of DONV (NC016997.1), CHOAV (NC017086.1), NOUV (EU159426.2), KPKV (KY320648.1), and ZIKV (LC002520.1) were synthesized by SYKMGENE Beijing and cloned into a pCDNA3.1 vector. RVPs were produced by co-transfection of 9 μg CprME expressing plasmid and 3 μg WNV replicon plasmid pWNVII-Rep-GFPZeo encoding a GFP reporter into 293T cells by calcium phosphate. After 12 h, the medium was replaced with low-glucose DMEM plus 2% fetal bovine serum. The RVPs were collected 72 h post-transfection and stored at −80 °C.

To test the secretion of RVPs, supernatant was spin by an ultra-centrifuge at 30,000 rpm for 2 h through a 20% sucrose cushion. The WNV replicon RNAs were isolated from the pellet and reverse-transcribed using Prime Script™ RT reagent Kit with gDNA Eraser (Takara, Japan). Quantitative PCRs were performed using a SYBR PremixEX Taq II (RT)-PCR kit (Takara, Japan) on a Thermo PIKOREAL 96 real-time PCR System. The following amplification program was used: incubation at 95°C for 30 s, followed by 40 cycles of 95°C for 5 s and 60°C for 20 s. Information collection and melt curve analysis were done following the instrument’s operation manual.

### Immunofluorescence assay

Virus-infected cells were fixed with 4% paraformaldehyde (PFA) for 30 min and permeabilized with 0.5% Triton X-100 for 10 min. Then cells were blocked with 3% bovine serum albumin in PBS for 1 h, and then cells were incubated with mouse mAb 4G2 and mAb J2 (Scicons, EU) at a dilution of 1:400 for 3 h to detect flavivirus E protein and double-strand RNA, respectively. After being washed three times with PBS, cells were incubated with secondary antibody Alexa Fluor 488 goat anti-mouse at a dilution of 1:400 for 1 h. Hoechst 33342 was added at 1 μg/ml to stain nuclei. The resulting fluorescence was detected by confocal microscopy (Zeiss LSM 710; Germany).

### Endocytosis assay

Huh7.5 cells were seeded in a 24-well plate. Twenty-four hours later, ZIKV and DONV were pre-bound to the cells on ice for 30 min at a multiplicity of infection (MOI) of 100. Then cells were cultured at 37°C for 20 min following by fixation with 4% PFA for 30 min at RT. Endocytosis of virions was detected by 4G2 antibody and imaged by confocal microscopy as mentioned above.

### Virus replication assay

DONV, ZIKV, or chimeric infectious clone plasmid was transfected into C6/36 or 293T cells by FuGENE 6 transfection reagent (Promega, USA). At 72 h post-transfection, cellular RNAs were isolated by Trizol reagent (Invitrogen, USA). RNAs were treated for 1 h by DNase I at 37°C and reverse-transcribed using a Prime ScriptTM RT reagent kit with gDNA Eraser (Takara, Japan). Quantitative PCR was performed as mentioned above. The detection limit of DONV by real-time PCR was 49.8 copies.

### Tobramycin RNA affinity chromatography

Tobramycin RNA affinity chromatography was adapted from Hartmuth *et al* (59). NHS-activated sepharose beads (BEAVER, China) were washed four times with 1 mM HCl and resuspended in coupling buffer (0.2 M NaHCO3, 0.5 M NaCl, 5 mM tobramycin, pH 8.3). Following overnight incubation at 4°C with head-to-tail rotation, beads were collected by magnetic frame and resuspended in blocking buffer (100 mM Tris-HCl, 150 mM NaCl, pH8.0). After 3 h incubation at 4°C, beads were washed 3 times and resuspended in PBS. RNAs in binding buffer (20 mM Tris, pH 7.0, 1 mM CaCl_2_, 1 mM MgCl_2_, 75 mM NaCl, 145 mM KCl, 0.1 mg/ml yeast tRNA, and 0.2 mM DTT) were heated to 95°C for 5 min, cooled to room temperature for refolding, and then incubated at 4°C for 2 h with head-to-tail rotation to combined with tobramycin beads. Beads were centrifuged and washed four times with washing buffer (20 mM Tris, pH7.0, 1 mM CaCl_2_, 1 mM MgCl_2_, 75 mM NaCl, 145 mM KCl, 0.5% NP-40, and 0.2 mM DTT). Before binding with the RNA beads, cell lysate was pre-cleared using tobramycin beads. After pre-clearing, beads and lysate were incubated for 2 h at 4°C. Beads were collected by magnetic frame and washed four times with washing buffer (20 mM Tris, pH 7.0, 1 mM CaCl_2_, 1 mM MgCl_2_, 75 mM NaCl, 145 mM KCl, and 0.5% NP-40) and then incubated for 5 min at room temperature with elution buffer (20 mM Tris, pH 7.0, 1 mM CaCl_2_, 3 mM MgCl_2_, 145 mM KCl, 10 mM tobramycin, and 0.2 mM DTT). Elutions were collected by centrifugation.

### Mass spectrometry

Eluted protein complexes from RNA affinity chromatography were digested with trypsin, and then tryptic peptides were subjected to LC-MS/MS on an Easy-Nlc 1000 coupled with Q Exactive Orbitrap instrument (Thermo Fisher Scientific, USA) under standard conditions. The search parameters allowed for fixed cysteine methylthiolation and variable methionine oxidation modifications, with a 10 ppm peptide mass tolerance, 0.05 Da fragment mass tolerance, and one missed tryptic cleavage. Raw spectral files were converted to mascot generic format using MSGUI, then searched against a database containing human proteins from UniProt.

### Protein expression and purification

The coding sequences of WTAP, SYNCRIP, and G3BP1 were cloned into the pET28a vector by homologous recombination. The recombinant proteins carrying a C-terminal His-tag were expressed in *E. coli* Rosetta (DE3) in the presence of 0.1 mM isopropyl-β-D-thiogalactopyranoside (IPTG) at 16°C for 20 h. The cell pellets were harvested by centrifugation at 6000 g for 15 min and then resuspended in binding buffer (50 mM Tris-Cl, pH 8.0, and 300 mM NaCl) containing a protease inhibitor cocktail. After high pressure homogenization, the supernatants were centrifuged at 12000 rpm for 30 min. The supernatants were purified through Ni-chelating affinity chromatography. The recombinant proteins were eluted by binding buffer containing 50, 100, and 200 mM imidazole. All purified proteins were dialyzed in binding buffer, split into single-use aliquots, and stored at −80°C.

### Electrophoretic mobility shift assay (EMSA)

To test protein-RNA interactions, RNA segments were first diluted in 0.5 X TE buffer and heated at 95°C for 2 min and then placed on ice immediately. A 5 X RNA folding buffer (250 mM Tris-Cl, pH 8.0, 500 mM NaCl, and 25 mM MgCl_2_) was added to the samples, and the RNAs were refolded at 37°C for 20 min. The binding reactions contained 50 nM RNA, 5 X EMSA buffer (200 mM Tris-HCl, pH 8.0, 300 mM NaCl, and 25 mM MgCl_2_), 0.05 mg/ml heparin sodium salt, and 7.5% glycerol and different amounts of proteins (0, 2.5, 5, and 10 ug) in a 20 ul volume. The reactions were incubated at 30°C for 30 min. Then 10 X gel loading solution was added, and the mixtures were separated by electrophoresis on 6% native PAGE gels running in 0.5 X TBE at 4°C to prevent the dissociation of RNA-protein complexes and RNA degradation. After running at 60 V for 4 h, the gels were stained with Gel Red nucleic acid gel stain for 30 min, and pictures were captured using Image lab software with a Universal Hood III instrument (Bio-Rad, USA).

## Acknowledgments

We thank Prof. Zhengfan Jiang (College of Life Sciences, Peking University) for providing Hela cell and its KO derivatives. We thank Dr. Zhe Lin (Institute of Zoology, CAS) for preparing Figure 5. We thank Dr. Yong Zhang (Institute of Zoology, CAS) for assistance of phylogenetic analysis. Funding: This project was funded by the National Key Plan for Scientific Research and Development of China (2016YFD0500303), National Natural Science Foundation of China, General Program (81871687), the National Science and Technology Major Project (2018ZX10101004), and Open Research Fund Program of State Key Laboratory of Integrated Pest Management (IPM1806).

---

**Supplementary Figure 1.**
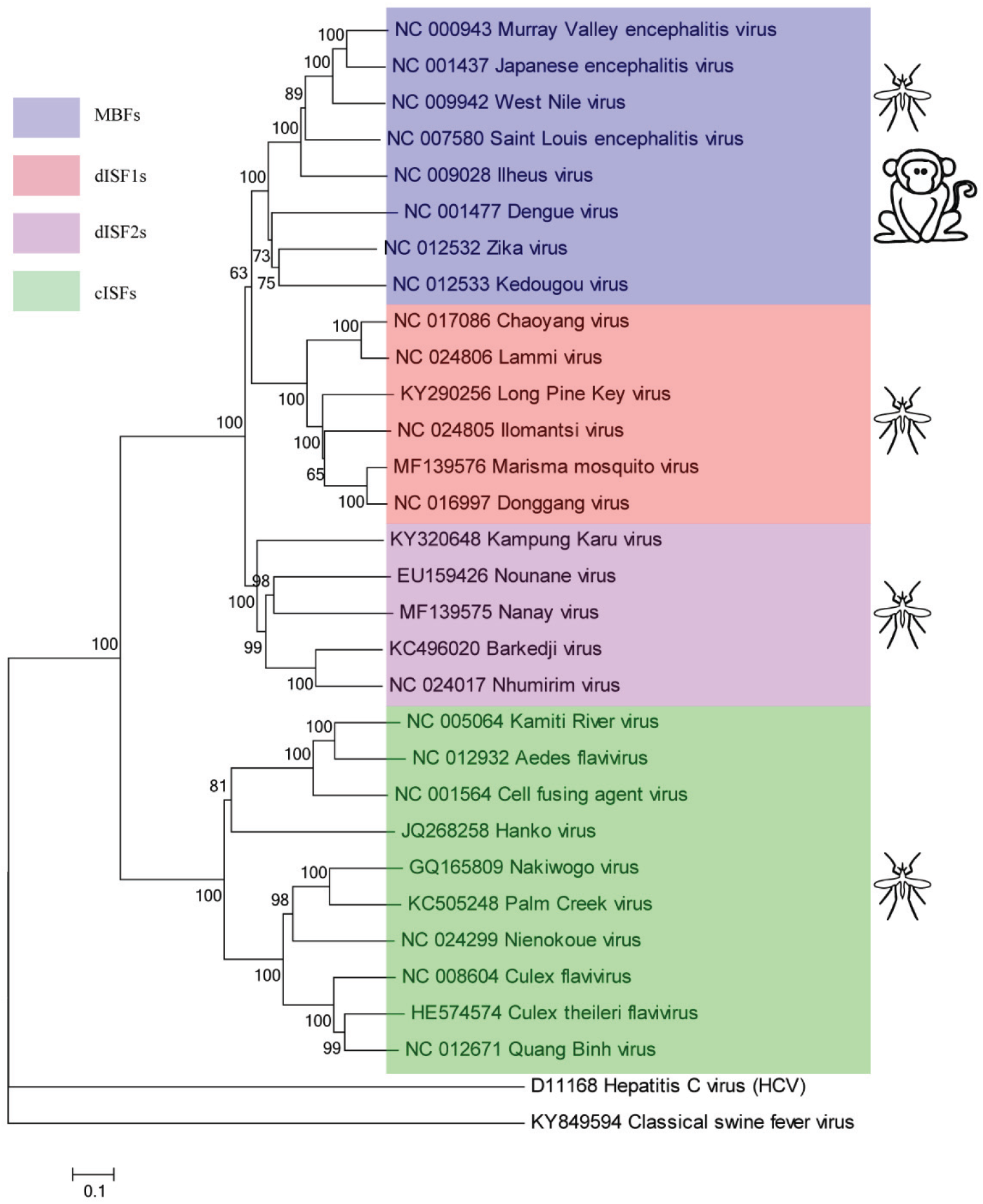
Phylogenetic tree of different groups of flaviviruses introducing Hepatitis C virus and classical swine fever virus as the outgroup. Complete polyprotein amino acid sequences were aligned and a maximum likelihood phylogenetic tree was reconstructed in MEGA7(51). The consensus tree inferred based on 500 replicates (52) is taken to represent the evolutionary history of the taxa, where the bootstrap values are shown along the branch. Highlighted in blue are mosquito-borne flaviviruses (MBFs), red are dual-host affiliated insect-specific flaviviruses 1 (dISF1s), purple are dual-host affiliated insect-specific flaviviruses 2 (dISF2s), and green are classical insect-specific flaviviruses (cISFs).

**Supplementary Figure 2.**
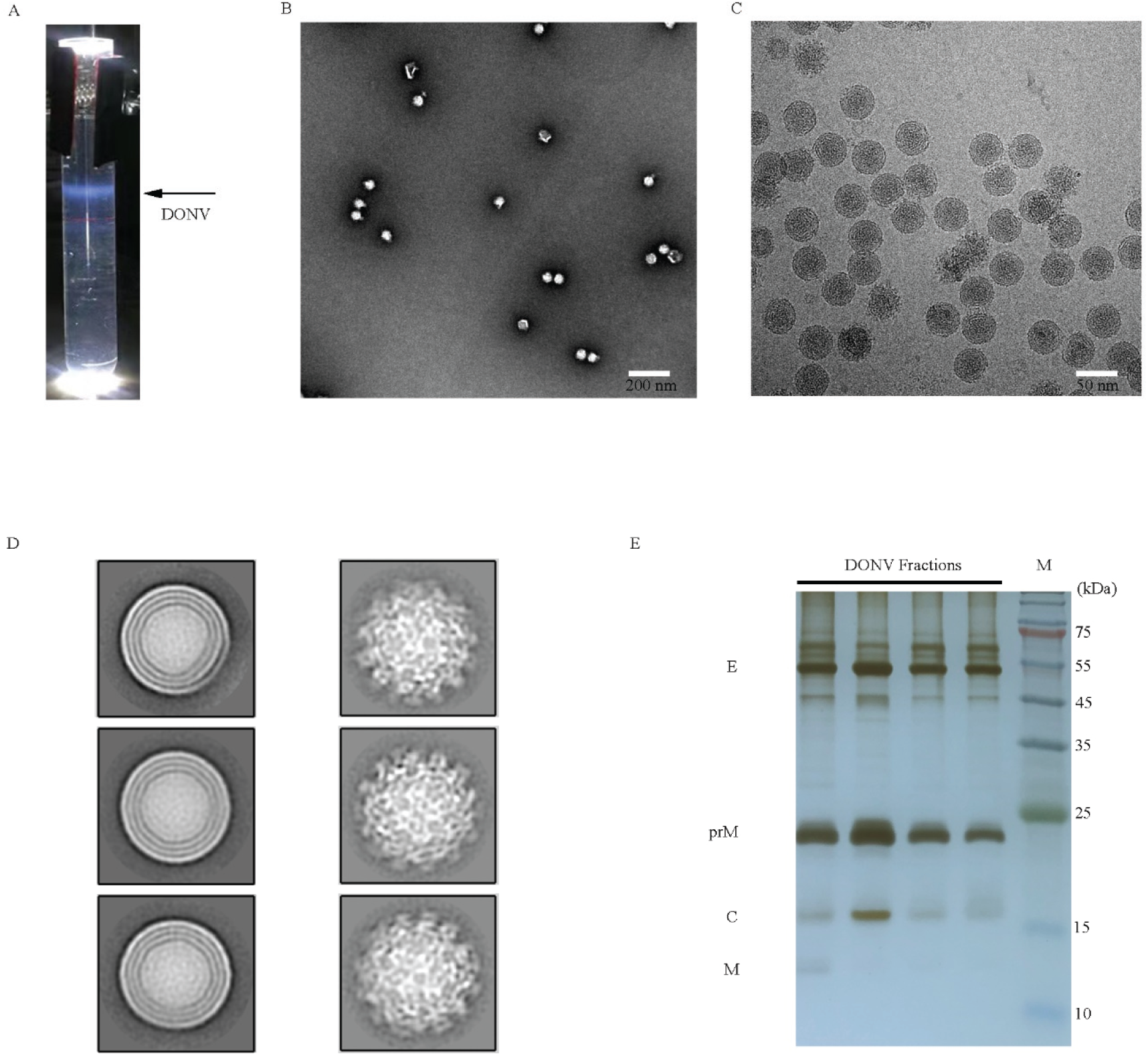
Purification and data collection of DONV. (A) Zonal ultracentrifugation of a 15–45% (w/v) sucrose density gradient at 30,000 rpm for 3 h was used to purify DONV from the harvest concentrate described in the Methods section. (B) The negative-stain images of DONV. (C) The cryo-EM micrograph of DONV. (D) Representative classes from 2D classification in RELION for DONV. (E) SDS-PAGE analysis of continuous sucrose gradient ultracentrifugation purified DONV virions followed by silver staining.

**Supplementary Figure 3.**
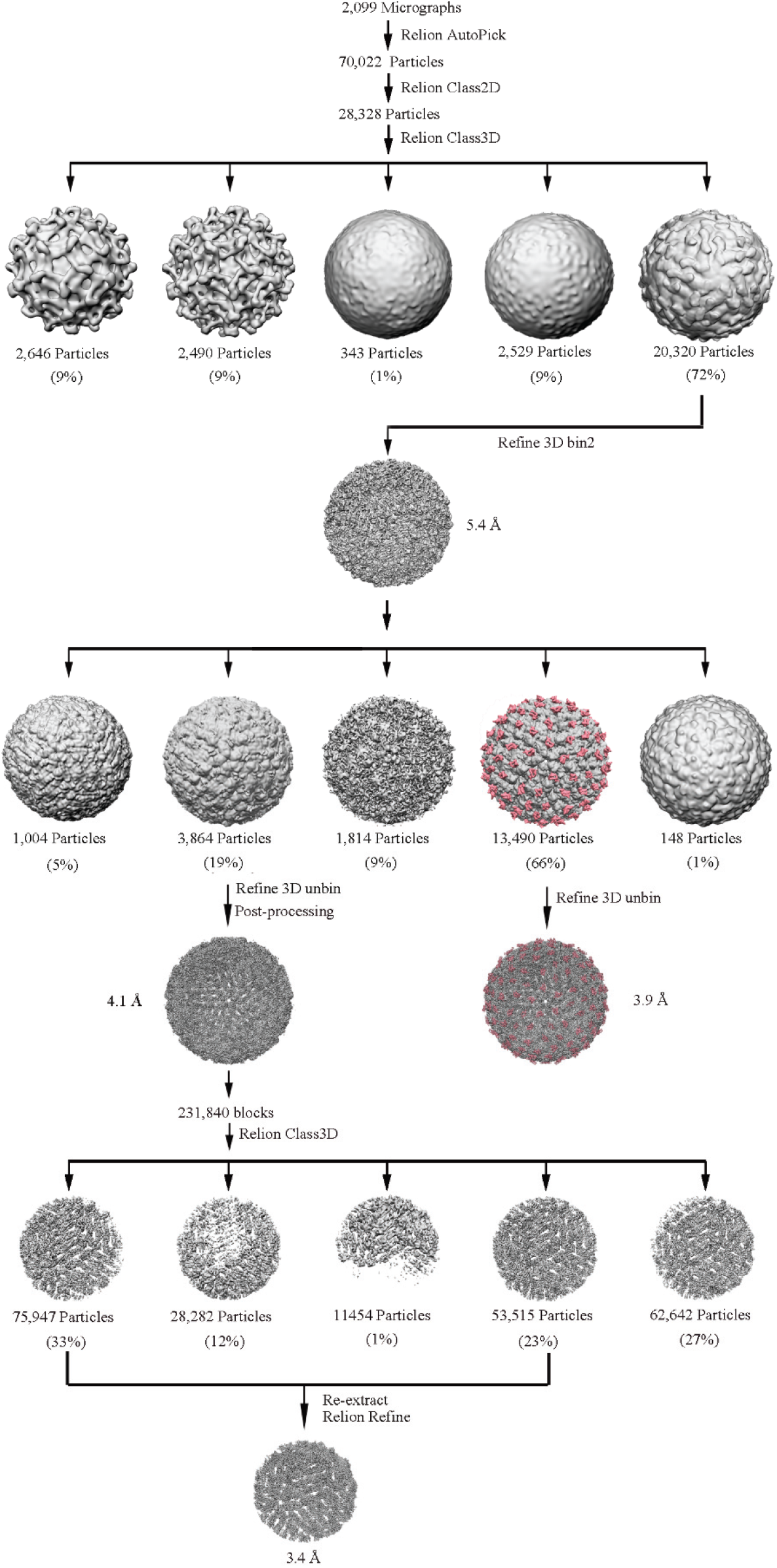
Flow-chart for Cryo-EM data processing. The data processing procedures for DONV. Details are given in the Methods section.

**Supplementary Figure 4.**
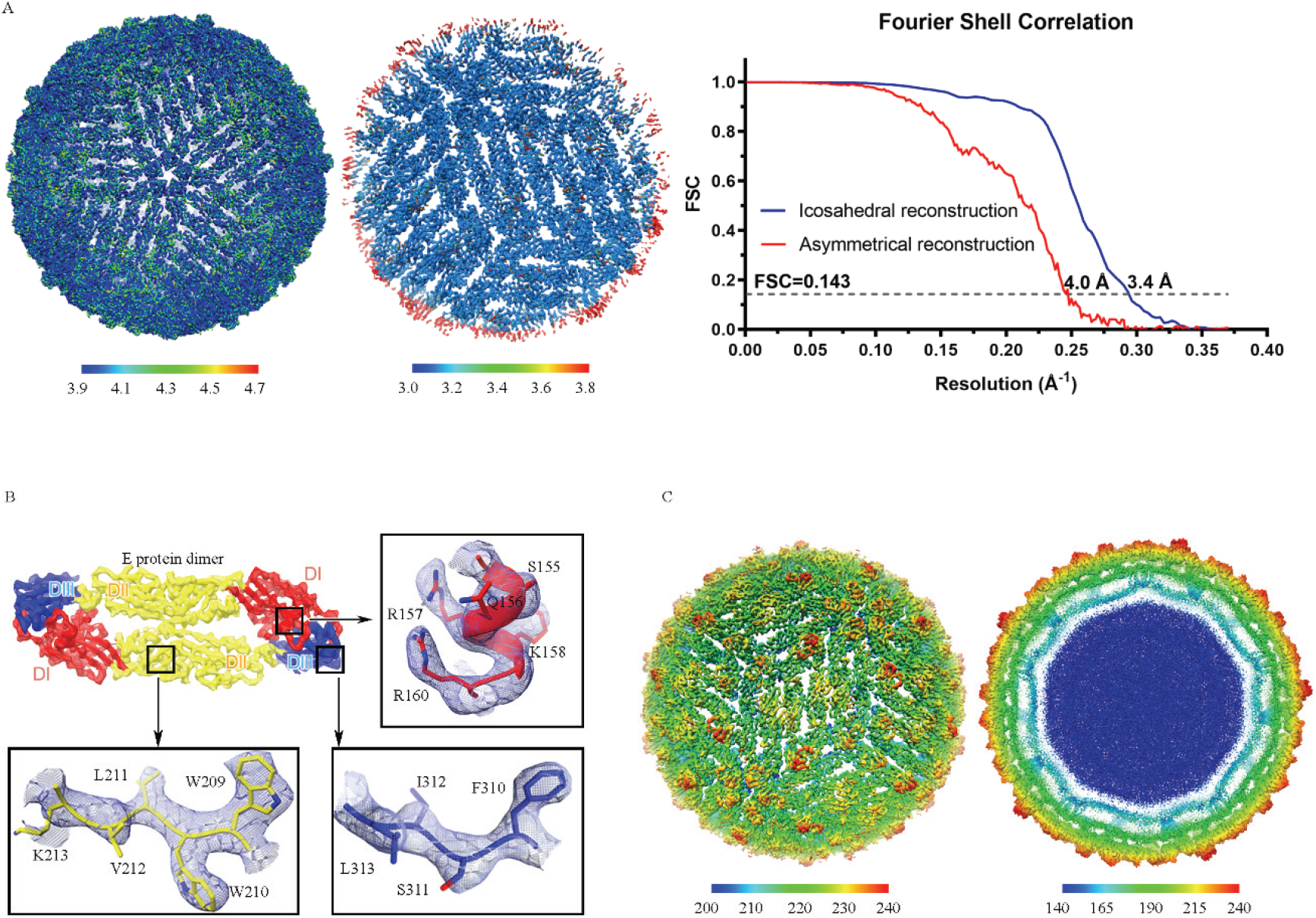
Assessment of resolution and electron density maps. (A) The map resolution assessment of icosahedral and asymmetrical reconstructions, showing resolutions distribution from 3.9 to 4.7 Å and 3.0 to 3.8 Å. The gold-standard FSC curves of icosahedral and asymmetrical reconstructions. The resolutions at FSC=0.143 are 4.0 Å and 3.4 Å. (B) Electron density maps for a section of the envelope protein DI, DII, and DIII domain. (C) Cryo-EM map of DONV viewed down an icosahedral twofold axis.

**Supplementary Figure 5.**
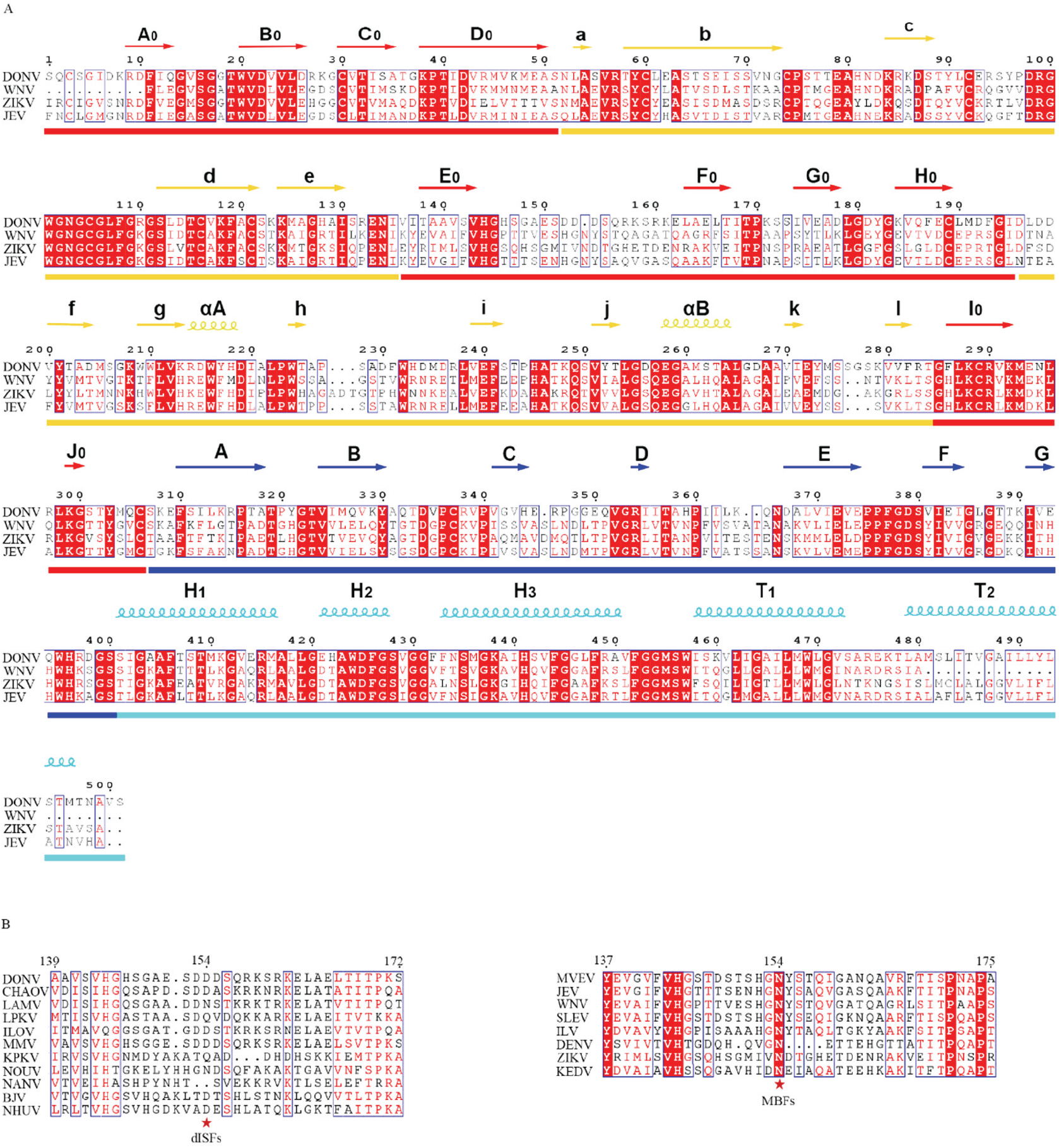
Sequence alignment of viruses shown in figure 1. (A) Alignment of E protein sequences of DONV with WNV, ZIKV, and JEV. DI, DII, DIII, and stem (H1, H2, H3, T1, and T2) are indicated below the sequences and shown in red, yellow, blue, and cyan, respectively. The secondary structures of DONV are labeled above the sequences. (B) Sequence alignment of E proteins glycosylation motif from representative dISFs and MBFs. Potential N-glycosylation site amin acid 153/154 is labeled with asterisk.

**Supplementary Figure 6.**
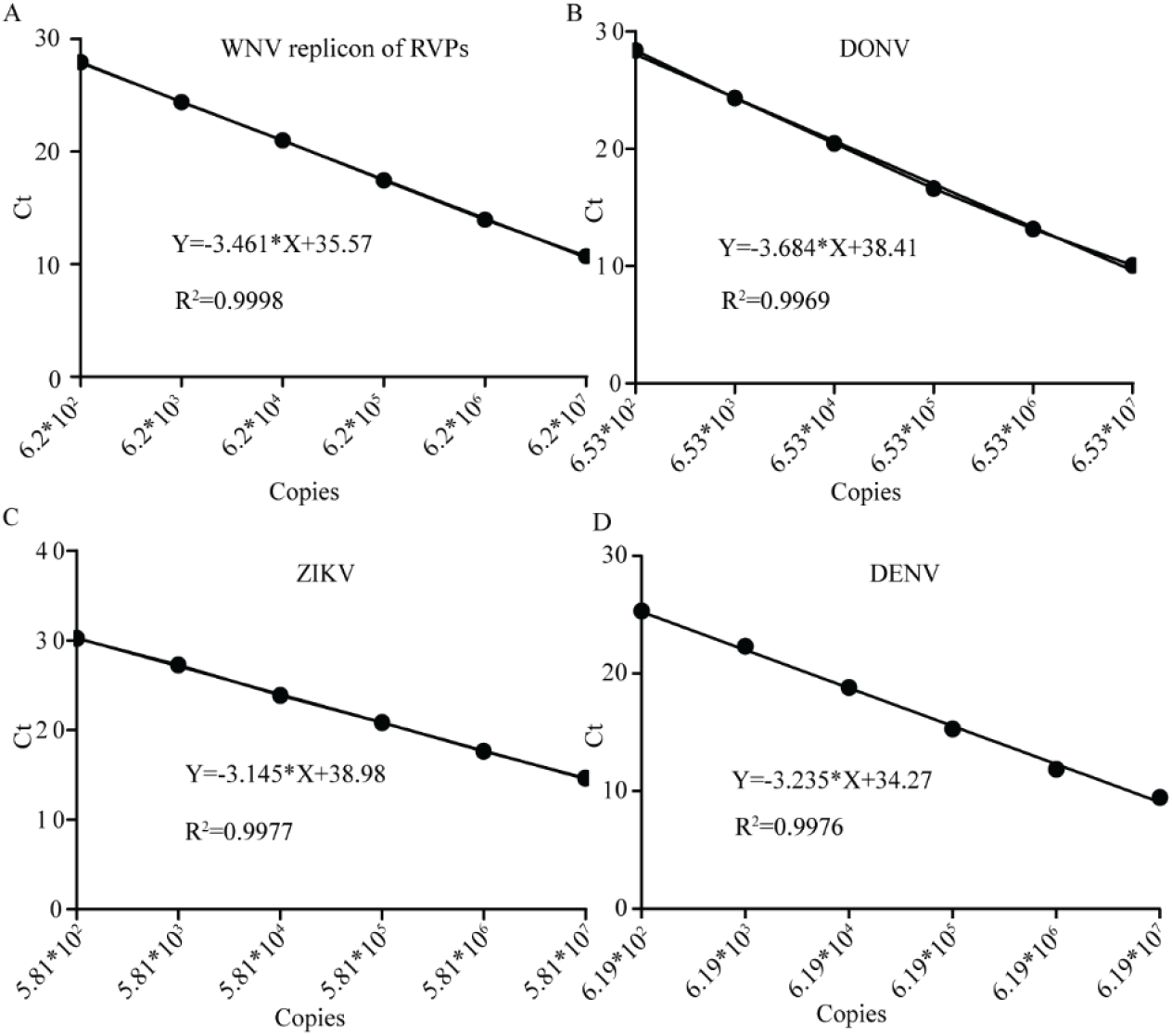
Standard curve for the real-time PCR assays for the detection of WNV replicon in the RVPs (A), DONV (B), ZIKV (C), and DENV (D). The sizes of the PCR products are 105 bp, 132 bp, 122 bp, and 143 bp for WNV replicon, DONV, ZIKV, and DENV, respectively. The correlation coefficients (R^2^) were 0.9998, 0.9969, 0.9977, and 0.9976 for WNV (A), DONV (B), ZIKV (C), and DENV (D), respectively. The linear dynamic ranges of both assays were higher than 6 logs, and all the threshold cycle (Ct) values in the following experiments fell into the dynamic ranges. The results represent two independent experiments.

**Supplementary Figure 7.**
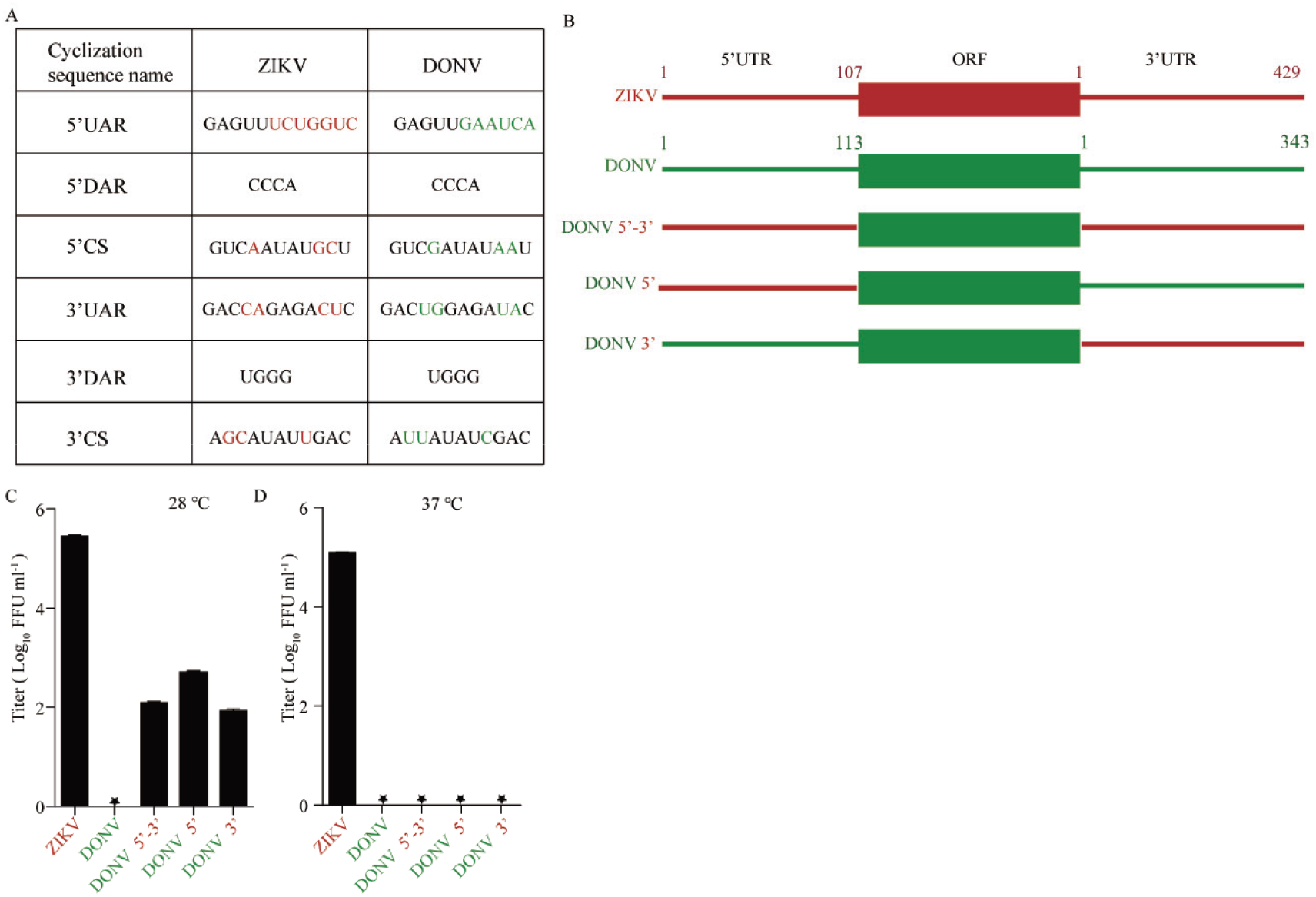
Release of DONV 5’, DONV 3’, and DONV 5’-3’ was partially rescued at 28 °C. (A) Cyclization sequences of ZIKV and DONV. (B) Schematic diagram of chimeric infectious clones. DONV 5’-3’, the UTRs of DONV replaced by those of ZIKV; DONV 5’, the 5’ UTR of DONV replaced by ZIKV; DONV 3’, the 3’ UTR of DONV replaced by ZIKV. (C and D) ZIKV, DONV, and chimeric infectious clones were transfected into 293T cells. The titers of viruses in the supernatants at 28 °C and 37 °C were tested by a focus-forming assay. n=3. Error bars indicate SD. These results represent three independent experiments.

**Supplementary Figure 8.**
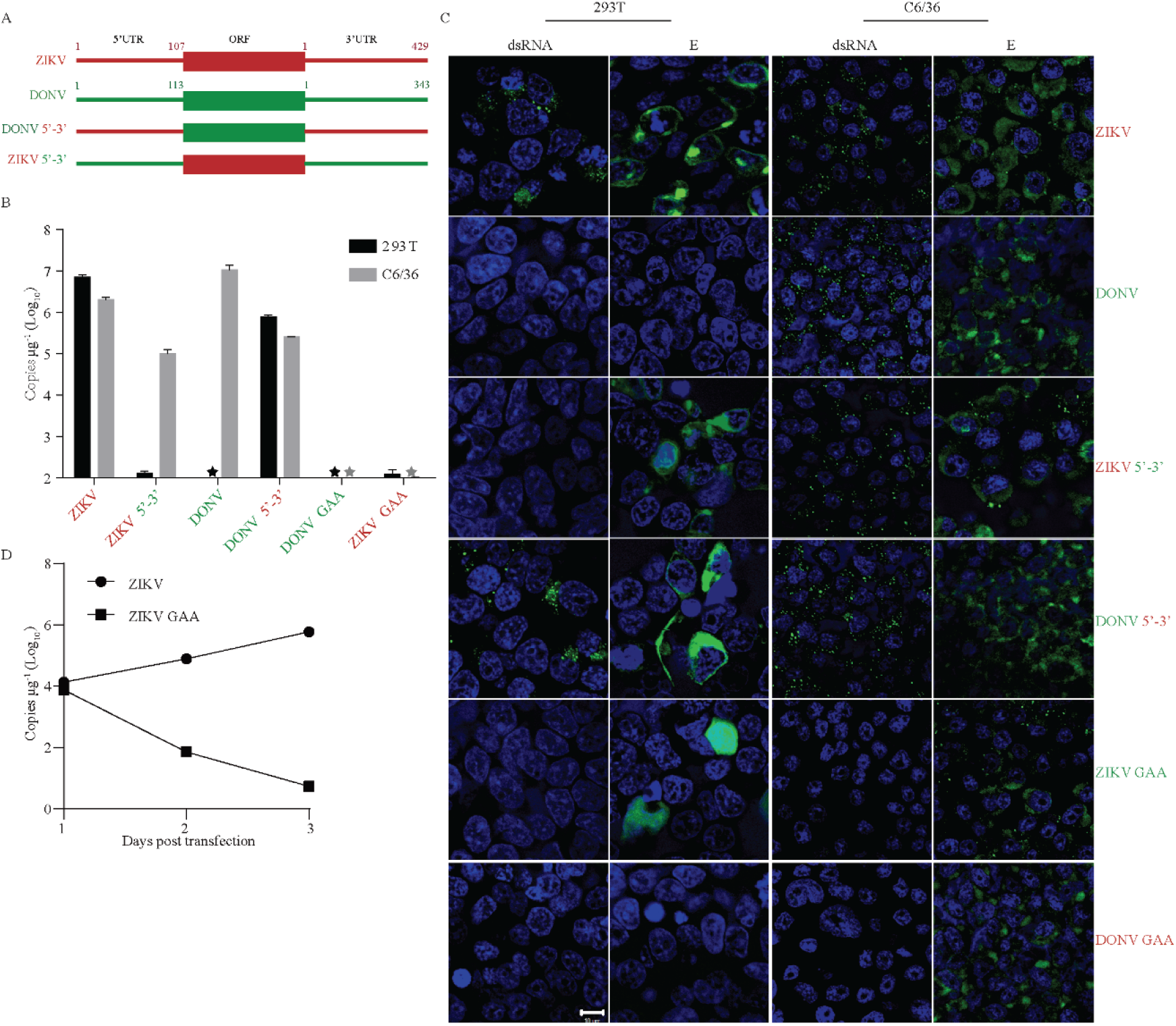
Replacement of ZIKV UTRs with those of DONV abolished ZIKV replication in vertebrate cells. (A) Schematic diagram of chimeric infectious clones. DONV 5’-3’, the UTRs of DONV replaced by those of ZIKV; ZIKV 5’-3’, the UTRs of ZIKV replaced by those of DONV. (B) Viral RNA replication of WT and chimeric viruses in vertebrate cells and C6/36 cells as measured by real-time PCR. n=3. Error bars indicate SD. (C) Immunofluorescence analysis of viral double-strand RNA by anti-dsRNA mAb and envelop protein E by 4G2 mAb of chimeric infectious clones in transfected human 293T cells and mosquito C6/36 cells. (D) The viral RNA levels of GAA mutants in 293T cells in (B) as measured at different time-points. WT and GAA ZIKV infectious clones were transfected into 293T cells and viral RNAs were measured by real-time PCR at indicated time-points. These results represent three independent experiments.

**Supplementary Figure 9.**
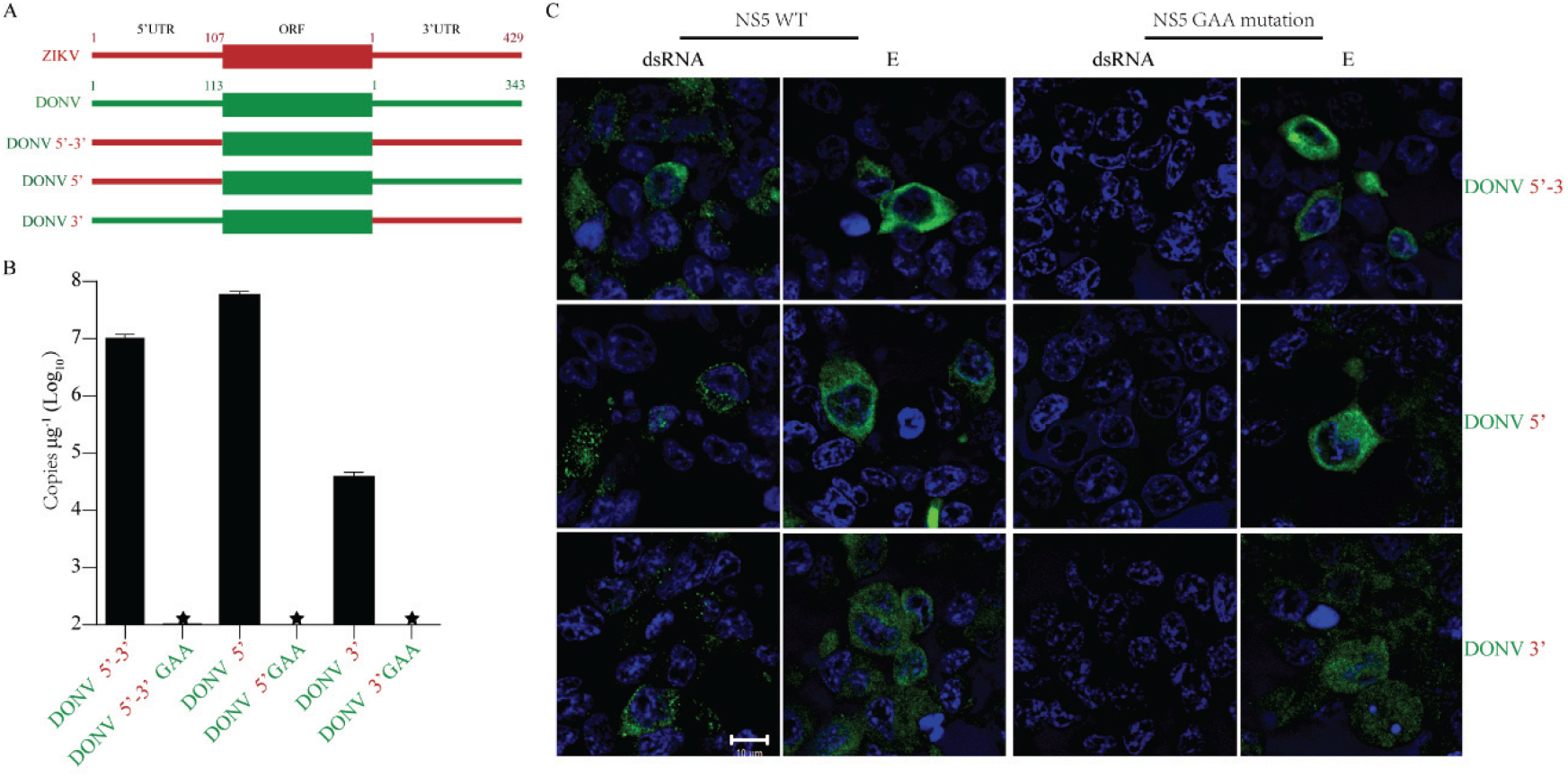
GAA mutation in the NS5 protein abolished the replication of DONV 5’, DONV 3’, and DONV 5’-3’ without affecting viral protein translation in 293T cells. (A) Schematic diagram of chimeric infectious clones. (B) 293T cells were transfected with the infectious clones of viruses with WT or GAA mutant NS5. Replication of viruses in 293T cell lysates were measured by real-time PCR 48 h later. n=3.Error bars indicate SD. (C) Double-strand RNA (dsRNA) and E protein were detected by immunofluorescence with mAb against dsRNA and 4G2 respectively (green). Nuclei were stained by Hoechst 33342 (blue). These results represent three independent experiments.

**Supplementary Figure 10.**
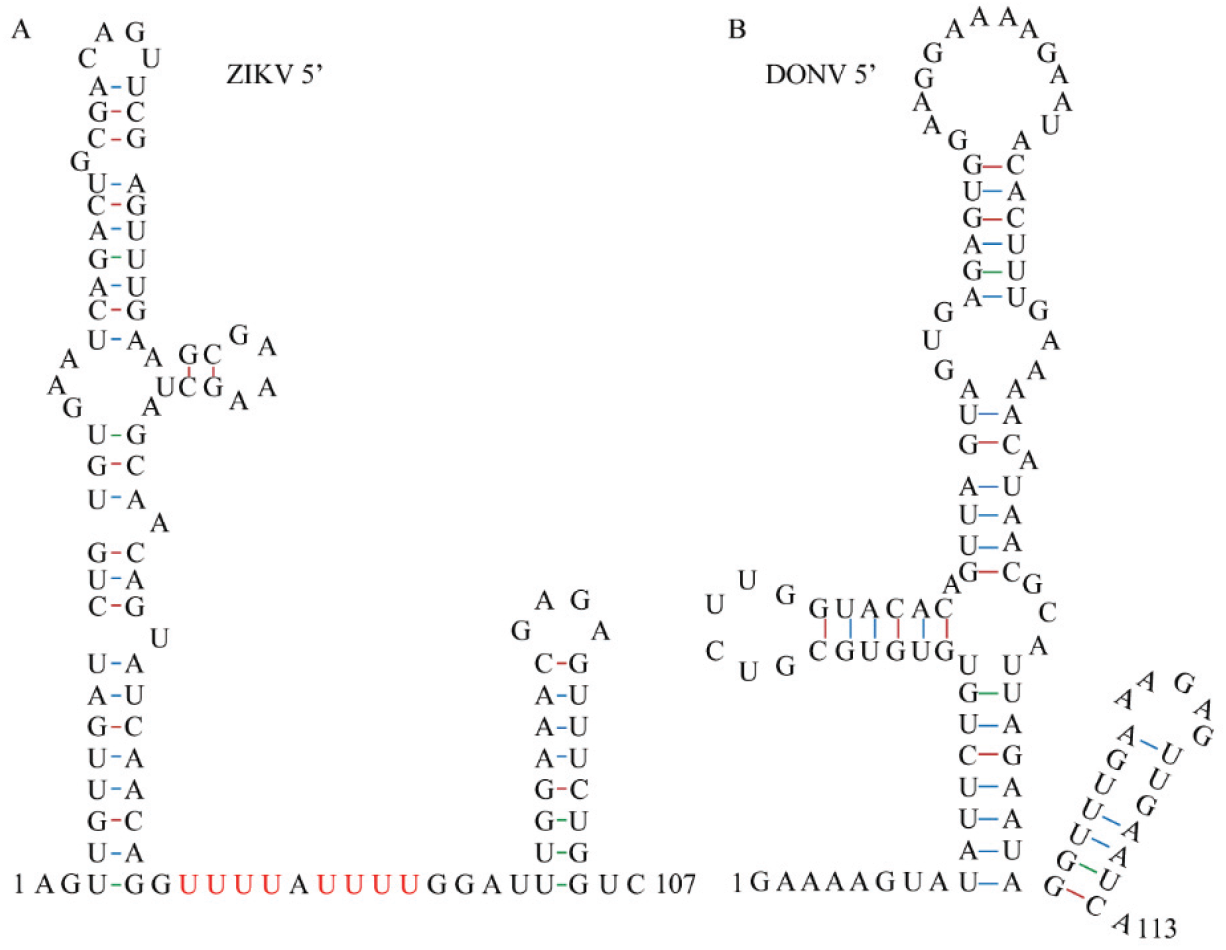
Predicted secondary structure of ZIKV (A) and DONV 5’ UTR (B) by RNAfold.

**Supplementary Figure 11.**
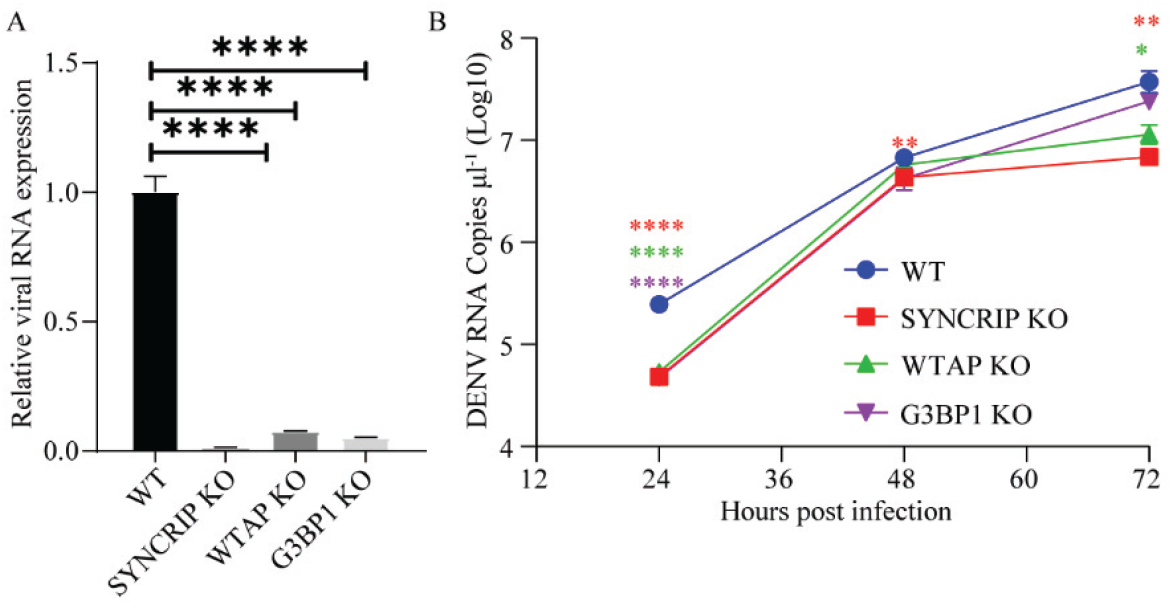
WTAP, SYNCRIP, and G3BP1 are critical for DENV replication in 293T cells. (A) Viral RNA replication in WT, WTAP-KO (knockout), G3BP1-KO, and SYNCRIP-KO cells. The cells were infected with DENV at an MOI of 0.1. RNAs were isolated by Trizol reagent 72 h post infection and tested by real-time PCR. The expression of Viral RNA was normalized to relative to β-actin. The value of WT was set as 1. (B) Viral RNA in the supernatants of WT, WTAP-KO, G3BP1-KO, and SYNCRIP-KO cells at indicated times were tested by real-time PCR. The data was analyzed by unpaired *t* test (A) and multiple *t* test (B). n=3. Error bars indicate SD. These results represent three independent experiments.

**Supplementary Figure 12.**
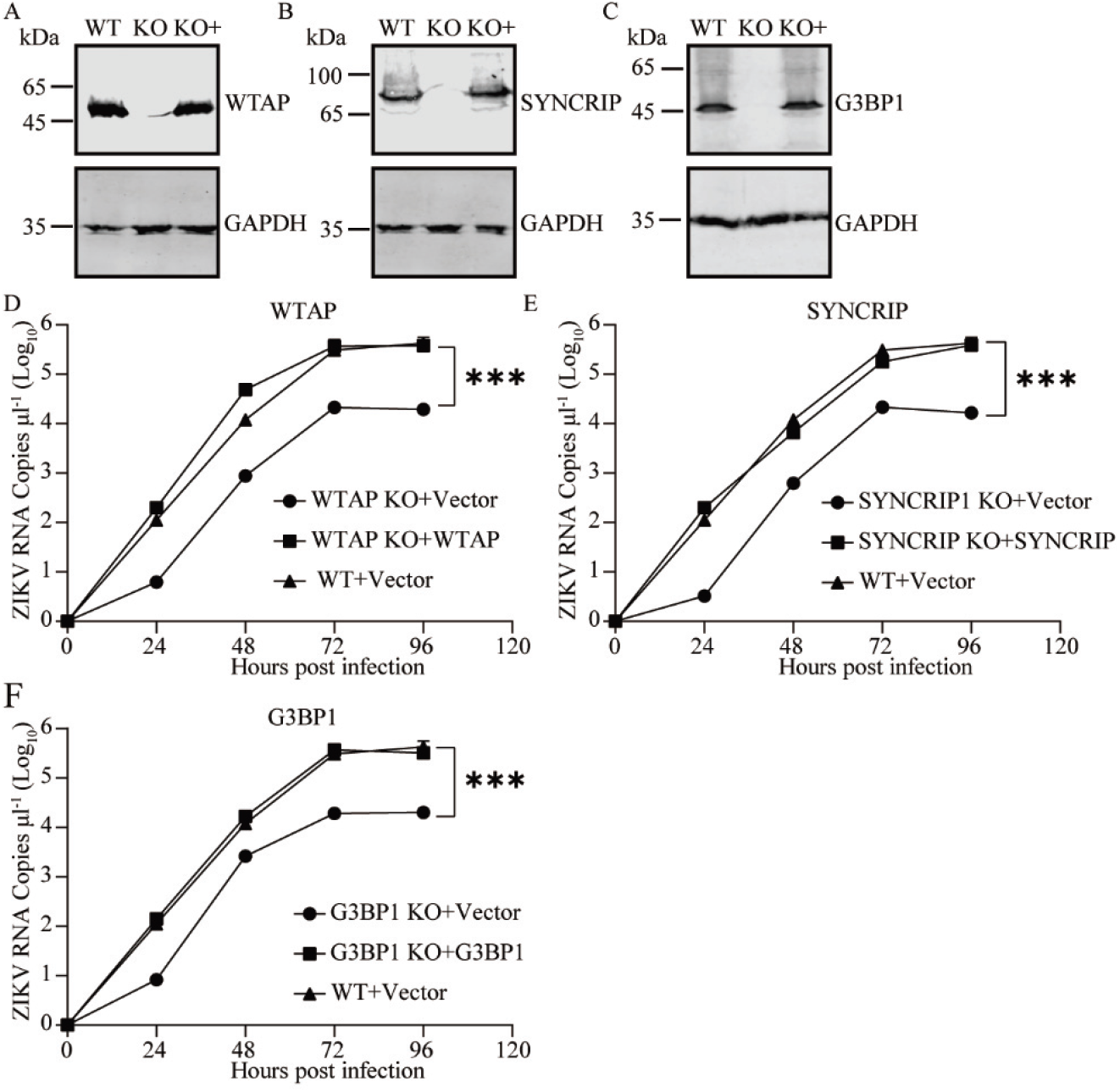
WTAP, SYNCRIP, and G3BP1 are critical for ZIKV replication in 293T cells. The expressions of WTAP (A), SYNCRIP (B), and G3BP1 (C) in WT, knockout (KO), and knockout+transduction (KO+) 293T cells were tested by Western blot with mAbs against indicated proteins. (D-F) WT, KO, and KO+ 293T cell lines were infected by ZIKV at an MOI of 0.1. Viral RNA in the supernatants was measured by real-time PCR at indicated times. The data was analyzed by multiple t test. n=3 and error bars indicate SD. The results represent three independent experiments.

**Supplementary Figure 13.**
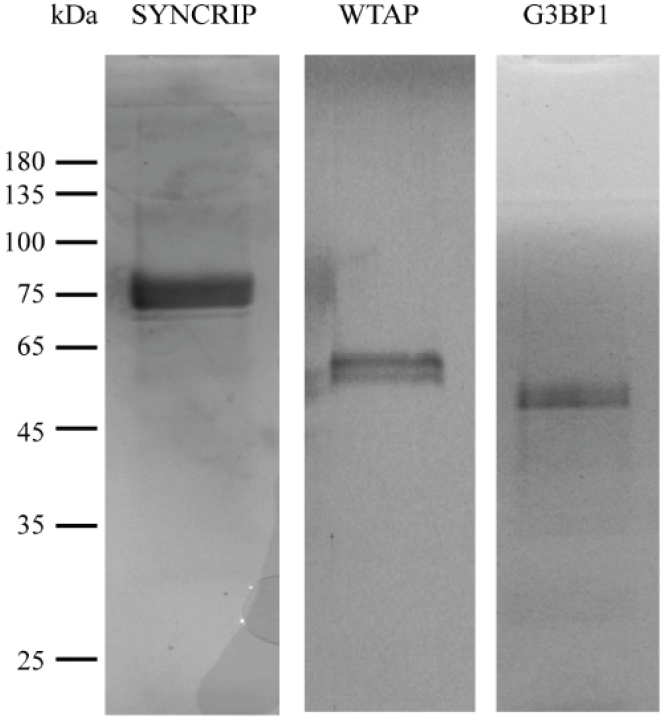
The Coomassie Blue staining of the purified proteins.

**Supplementary Figure 14.**
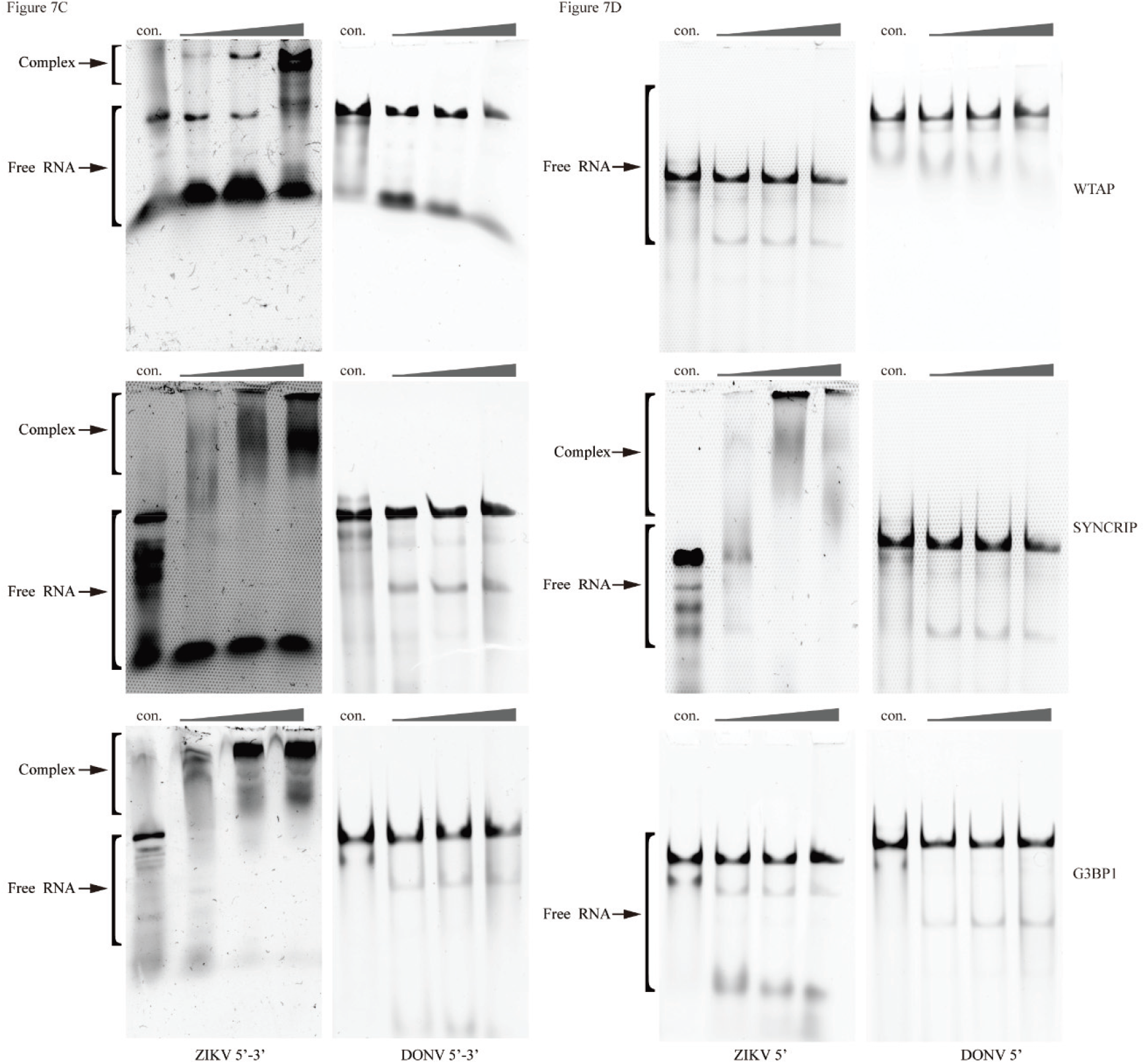
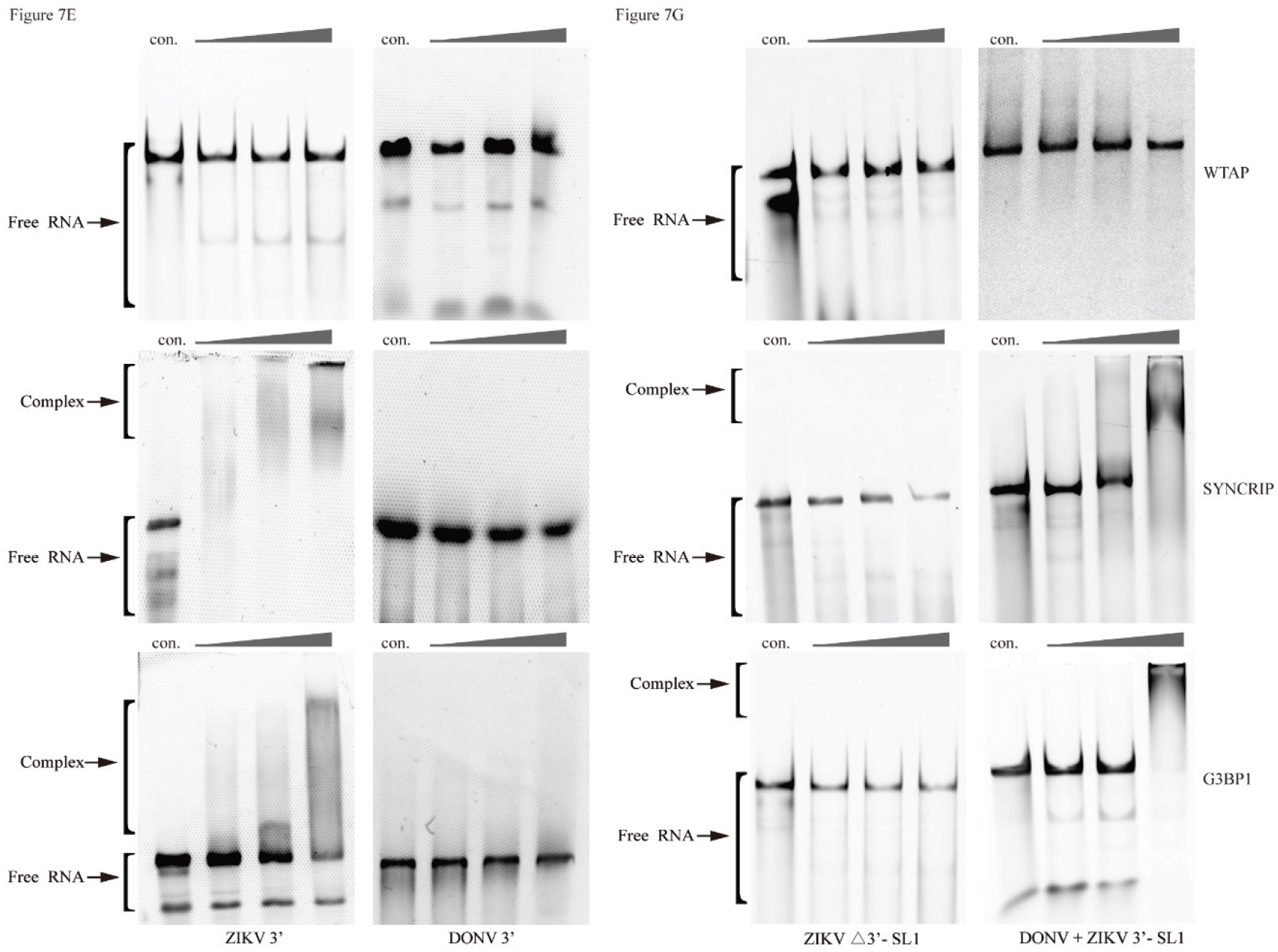
Original images of EMSA in Figure 7. Panels are labeled according to Figure 7.

**Supplementary Table 1.**
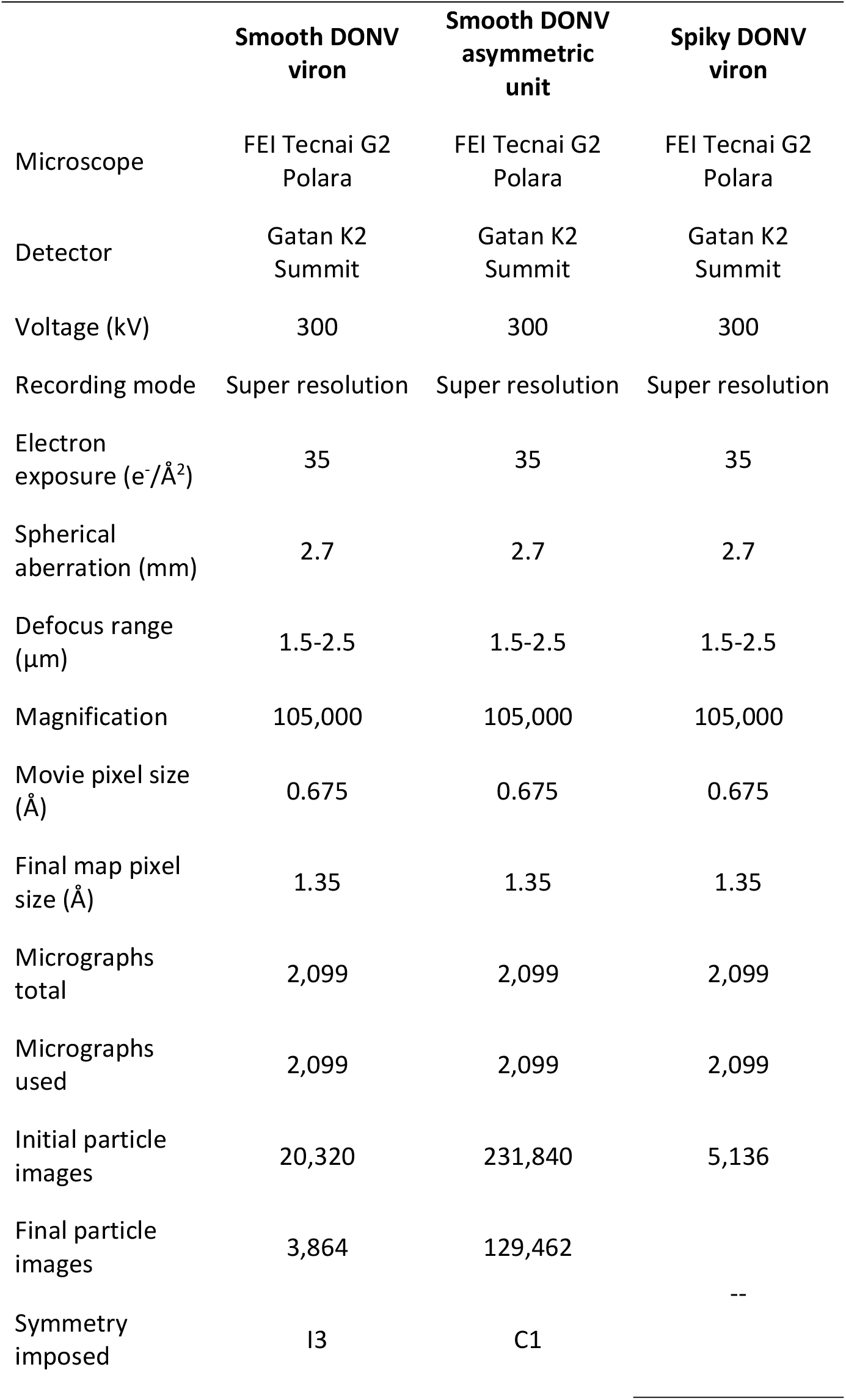
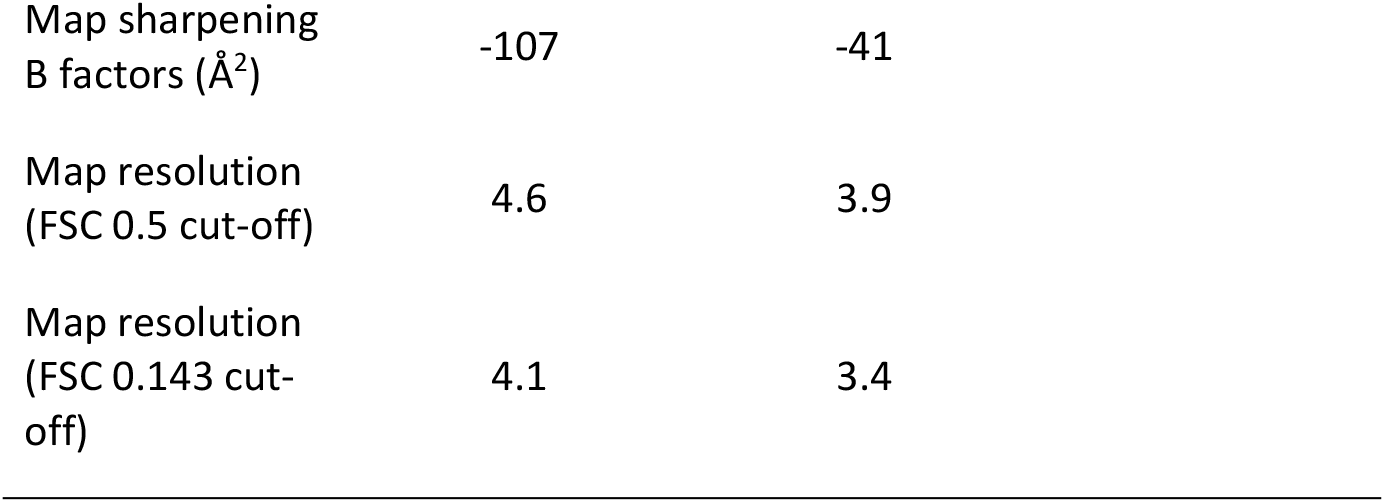
Cryo-EM data collection and processing statistics.

|  | Smooth DONV<br>vion | Smooth DONV<br>asymmetric<br>unit | Spiky DONV<br>vion |
| --- | --- | --- | --- |
| Microscope | FEI Tecnai G2<br>Polara | FEI Tecnai G2<br>Polara | FEI Tecnai G2<br>Polara |
| Detector | Gatan K2<br>Summit | Gatan K2<br>Summit | Gatan K2<br>Summit |
| Voltage (kV) | 300 | 300 | 300 |
| Recording mode | Super resolution | Super resolution | Super resolution |
| Electron<br>exposure (e<sup>-</sup>/Å<sup>2</sup>) | 35 | 35 | 35 |
| Spherical<br>aberration (mm) | 2.7 | 2.7 | 2.7 |
| Defocus range<br>(μm) | 1.5-2.5 | 1.5-2.5 | 1.5-2.5 |
| Magnification | 105,000 | 105,000 | 105,000 |
| Movie pixel size<br>(Å) | 0.675 | 0.675 | 0.675 |
| Final map pixel<br>size (Å) | 1.35 | 1.35 | 1.35 |
| Micrographs<br>total | 2,099 | 2,099 | 2,099 |
| Micrographs<br>used | 2,099 | 2,099 | 2,099 |
| Initial particle<br>images | 20,320 | 231,840 | 5,136 |
| Final particle<br>images | 3,864 | 129,462 | -- |
| Symmetry<br>imposed | I3 | C1 |  |

|  |  |  |
| --- | --- | --- |
| Map sharpening<br>B factors ($\text{\AA}^2$) | -107 | -41 |
| Map resolution<br>(FSC 0.5 cut-off) | 4.6 | 3.9 |
| Map resolution<br>(FSC 0.143 cut-off) | 4.1 | 3.4 |

**Supplementary Table 2.**
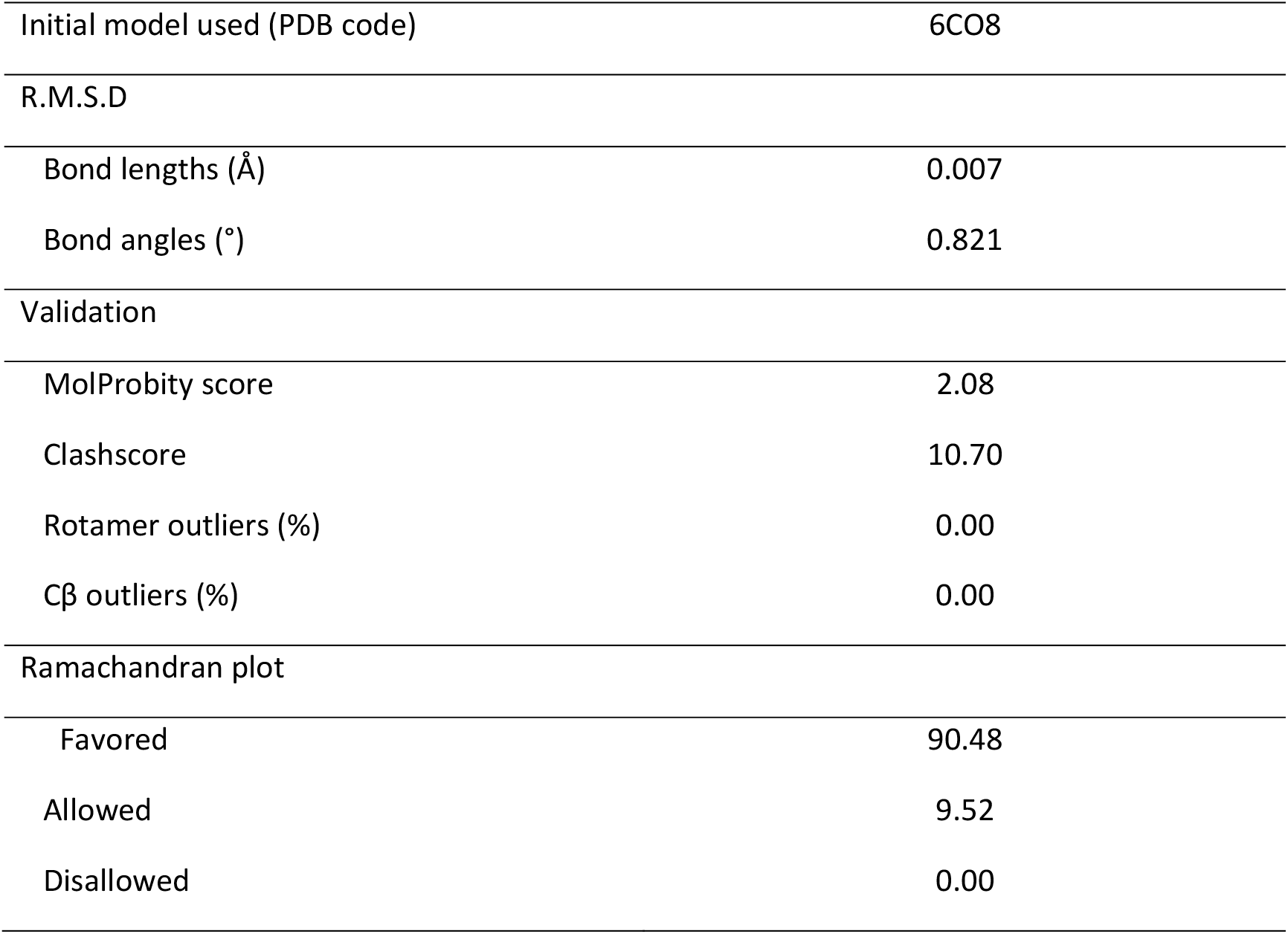
Model refinement statistics.

## References

1. Blitvich BJ & Firth AE (2015) Insect-specific flaviviruses: a systematic review of their discovery, host range, mode of transmission, superinfection exclusion potential and genomic organization. Viruses 7(4):1927–1959.

2. Gould EA & Solomon T (2008) Pathogenic flaviviruses. Lancet 371(9611):500–509.

3. Guzman H, et al. (2018) Characterization of Three New Insect-Specific Flaviviruses: Their Relationship to the Mosquito-Borne Flavivirus Pathogens. Am J Trop Med Hyg 98(2):410–419.

4. Calisher CH & Higgs S (2018) The Discovery of Arthropod-Specific Viruses in Hematophagous Arthropods: An Open Door to Understanding the Mechanisms of Arbovirus and Arthropod Evolution? Annu Rev Entomol 63:87–103.

5. Harrison JJ, et al. (2020) Antigenic Characterization of New Lineage II Insect-Specific Flaviviruses in Australian Mosquitoes and Identification of Host Restriction Factors. mSphere 5(3).

6. Tangudu CS, et al. (2021) Chimeric Zika viruses containing structural protein genes of insect-specific flaviviruses cannot replicate in vertebrate cells due to entry and post-translational restrictions. Virol. 559:30–39.

7. Wang ZSA, S.Y.; Wang, Y.; Han, Y.; Guo, J.Q. (2009) A new virus of Flavivirus: Chaoyang virus isolated in Liaoning province. Chin. Pub. Health 25:769–772.

8. Junglen S, et al. (2009) A new flavivirus and a new vector: characterization of a novel flavivirus isolated from uranotaenia mosquitoes from a tropical rain forest. J.Virol. 83(9):4462–4468.

9. Huhtamo E, et al. (2014) Novel flaviviruses from mosquitoes: mosquito-specific evolutionary lineages within the phylogenetic group of mosquito-borne flaviviruses. Virol. 464-465:320–329.

10. Huhtamo E, et al. (2009) Characterization of a novel flavivirus from mosquitoes in northern europe that is related to mosquito-borne flaviviruses of the tropics. J.Virol. 83(18):9532–9540.

11. McLean BJ, et al. (2020) The Insect-Specific Parramatta River Virus Is Vertically Transmitted by Aedes vigilax Mosquitoes and Suppresses Replication of Pathogenic Flaviviruses In Vitro. Vector borne and zoonotic diseases.

12. Goenaga S, et al. (2015) Potential for Co-Infection of a Mosquito-Specific Flavivirus, Nhumirim Virus, to Block West Nile Virus Transmission in Mosquitoes. Viruses 7(11):5801–5812.

13. Liu ZY, et al. (2016) Viral RNA switch mediates the dynamic control of flavivirus replicase recruitment by genome cyclization. Elife 5.

14. Lindenbach BD, Thiel H-J, & Rice CM (2007) Flaviviruses: the viruses and their replication. Fields’ Virology, eds Knipe DM & Howley PM (Lippincott, Williams and Wilkins, Philadelphia), Fifth Ed, pp 1101–1152.

15. Lorenz IC, Allison SL, Heinz FX, & Helenius A (2002) Folding and dimerization of tick-borne encephalitis virus envelope proteins prM and E in the endoplasmic reticulum. J.Virol. 76(11):5480–5491.

16. Zhang Y, et al. (2003) Structures of immature flavivirus particles. EMBO J. 22(11):2604–2613.

17. Bressanelli S, et al. (2004) Structure of a flavivirus envelope glycoprotein in its low-pH-induced membrane fusion conformation. EMBO J. 23(4):728–738.

18. Pokidysheva E, et al. (2006) Cryo-EM reconstruction of dengue virus in complex with the carbohydrate recognition domain of DC-SIGN. Cell 124(3):485–493.

19. Miller JL, et al. (2008) The mannose receptor mediates dengue virus infection of macrophages. PLoS Pathog. 4(2):e17.

20. Liu Y, et al. (2015) The roles of direct recognition by animal lectins in antiviral immunity and viral pathogenesis. Molecules 20(2):2272–2295.

21. Qiu X, et al. (2018) Structural basis for neutralization of Japanese encephalitis virus by two potent therapeutic antibodies. Nat Microbiol 3(3):287–294.

22. Wang X, et al. (2017) Near-atomic structure of Japanese encephalitis virus reveals critical determinants of virulence and stability. Nat Commun 8(1):14.

23. Matthew D. Therkelsen, et al. (2018) Flaviviruses have imperfect icosahedral symmetry PNAS 115(45):11608–11612.

24. Natalee D. Newton, et al. (2021) The structure of an infectious immature flavivirus redefines viral architecture and maturation. Sci. Adv. 7:eabe4507

25. Dong H, et al. (2021) Structural and molecular basis for foot-and-mouth disease virus neutralization by two potent protective antibodies. Protein Cell.

26. Nan Wang, et al. (2019) Architecture of African swine fever virus and implications for viral assembly. Science 366:640–644.

27. Fuzik T, et al. (2018) Structure of tick-borne encephalitis virus and its neutralization by a monoclonal antibody. Nat Commun 9(1):436.

28. Fibriansah G, et al. (2015) A highly potent human antibody neutralizes dengue virus serotype 3 by binding across three surface proteins. Nat Commun 6:6341.

29. Zhao H, et al. (2016) Structural Basis of Zika Virus-Specific Antibody Protection. Cell 166(4):1016–1027.

30. Wen D, et al. (2018) N-glycosylation of Viral E Protein Is the Determinant for Vector Midgut Invasion by Flaviviruses. mBio 9(1).

31. Ansarah-Sobrinho C, Nelson S, Jost CA, Whitehead SS, & Pierson TC (2008) Temperature-dependent production of pseudoinfectious dengue reporter virus particles by complementation. Virol. 381(1):67–74.

32. Atieh T, et al. (2017) New reverse genetics and transfection methods to rescue arboviruses in mosquito cells. Scientific reports 7(1):13983.

33. Lott WB & Doran MR (2013) Do RNA viruses require genome cyclisation for replication? Trends Biochem.Sci. 38(7):350–355.

34. Lodeiro MF, Filomatori CV, & Gamarnik AV (2009) Structural and functional studies of the promoter element for dengue virus RNA replication. J.Virol. 83(2):993–1008.

35. Zhang Y, et al. (2019) Long non-coding subgenomic flavivirus RNAs have extended 3D structures and are flexible in solution. EMBO Rep 20(11):e47016.

36. Lan J, et al. (2020) Structure of the SARS-CoV-2 spike receptor-binding domain bound to the ACE2 receptor. Nature 581(7807):215–220.

37. Shi M, et al. (2016) Redefining the invertebrate RNA virosphere. Nature.

38. Shi M, et al. (2016) Divergent Viruses Discovered in Arthropods and Vertebrates Revise the Evolutionary History of the Flaviviridae and Related Viruses. J.Virol. 90(2):659–669.

39. Piyasena TBH, et al. (2017) Infectious DNAs derived from insect-specific flavivirus genomes enable identification of pre- and post-entry host restrictions in vertebrate cells. Scientific reports 7(1):2940.

40. Junglen S, et al. (2017) Host Range Restriction of Insect-Specific Flaviviruses Occurs at Several Levels of the Viral Life Cycle. mSphere 2(1).

41. Auguste AJ, et al. (2021) Isolation of a novel insect-specific flavivirus with immunomodulatory effects in vertebrate systems. Virology 562:50–62.

42. Colmant AMG, et al. (2021) Insect-Specific Flavivirus Replication in Mammalian Cells Is Inhibited by Physiological Temperature and the Zinc-Finger Antiviral Protein. Viruses 13(4).

43. Colmant AMG, et al. (2017) A New Clade of Insect-Specific Flaviviruses from Australian Anopheles Mosquitoes Displays Species-Specific Host Restriction. mSphere 2(4).

44. Pijlman GP, et al. (2008) A highly structured, nuclease-resistant, noncoding RNA produced by flaviviruses is required for pathogenicity. Cell host & microbe 4(6):579–591.

45. Pallares HM, et al. (2020) Zika Virus Subgenomic Flavivirus RNA Generation Requires Cooperativity between Duplicated RNA Structures That Are Essential for Productive Infection in Human Cells. J Virol 94(18).

46. Lingling Zeng, Barry Falgout, & Markoff AL (1998) Identification of Specific Nucleotide Sequences within the Conserved 39-SL in the Dengue Type 2 Virus Genome Required for Replication. Journal of Virology 72(9):7510–7522.

47. Ngo KA, Rose JT, Kramer LD, & Ciota AT (2019) Adaptation of Rabensburg virus (RBGV) to vertebrate hosts by experimental evolution. Virol. 528:30–36.

48. Colmant AMG, et al. (2020) NS4/5 mutations enhance flavivirus Bamaga virus infectivity and pathogenicity in vitro and in vivo. PLoS Negl Trop Dis 14(3):e0008166.

49. Bidet K, Dadlani D, & Garcia-Blanco MA (2014) G3BP1, G3BP2 and CAPRIN1 are required for translation of interferon stimulated mRNAs and are targeted by a dengue virus non-coding RNA. PLoS Pathog. 10(7):e1004242.

50. Little NA, Hastie ND, & Davies RC (2000) Identification of WTAP, a novel Wilms’ tumour 1-associating protein. Hum Mol Genet 9(15):2231–2239.

51. Liu HM, et al. (2009) SYNCRIP (synaptotagmin-binding, cytoplasmic RNA-interacting protein) is a host factor involved in hepatitis C virus RNA replication. Virol. 386(2):249–256.

52. Crill WD & Chang GJ (2004) Localization and characterization of flavivirus envelope glycoprotein cross-reactive epitopes. J.Virol. 78(24):13975–13986.

53. Mastronarde DN (2005) Automated electron microscope tomography using robust prediction of specimen movements. J Struct Biol 152(1):36–51.

54. Li X, et al. (2013) Electron counting and beam-induced motion correction enable near-atomic-resolution single-particle cryo-EM. Nat Methods 10(6):584–590.

55. Scheres SH (2012) RELION: implementation of a Bayesian approach to cryo-EM structure determination. J Struct Biol 180(3):519–530.

56. Yang Y, et al. (2020) Architecture of the herpesvirus genome-packaging complex and implications for DNA translocation. Protein & cell 11(5):339–351.

57. Pettersen EF, et al. (2004) UCSF Chimera--a visualization system for exploratory research and analysis. J Comput Chem 25(13):1605–1612.

58. Afonine PV, et al. (2012) Towards automated crystallographic structure refinement with phenix.refine. Acta Crystallogr D Biol Crystallogr 68(Pt 4):352–367.

59. Hartmuth K, Vornlocher HP, & Lührmann R (2004) Tobramycin affinity tag purification of spliceosomes. Methods in molecular biology (Clifton, N.J.) 257:47–64.

